# Higher Angiotensin I Converting Enzyme 2 (ACE2) levels in the brain of individuals with Alzheimer’s disease

**DOI:** 10.1101/2023.01.17.524254

**Authors:** Reveret Louise, Leclerc Manon, Emond Vincent, Loiselle Andréanne, Bourassa Philippe, Tremblay Cyntia, David A Bennett, Hébert Sébastien, Calon Frédéric

**Author notes:** <u>Corresponding author:</u> Frédéric Calon, Centre de Recherche du CHUL (CHUQ) 2705, Boulevard Laurier, Room T2-05 Québec, QC, G1V 4G2, Canada.

## Abstract

The severe acute respiratory syndrome coronavirus 2 (SARS-CoV-2) is a major cause of death in the elderly. Cognitive decline due to Alzheimer’s disease (AD) is frequent in the geriatric population disproportionately affected by the COVID-19 pandemic. Interestingly, central nervous system (CNS) manifestations have been reported in SARS-CoV-2-infected patients. In this study, we investigated the levels of Angiotensin I Converting Enzyme 2 (ACE2), the main entry receptor of SARS-COV-2 in cells, in *postmortem* parietal cortex samples from two independent AD cohorts, totalling 142 persons. Higher concentrations of ACE2 protein and mRNA were found in individuals with a neuropathological diagnosis of AD compared to age-matched healthy control subjects. Brain levels of soluble ACE2 were inversely associated with cognitive scores (p = 0.02), markers of pericytes (PDGFRβ, p=0.02 and ANPEP, p = 0.007) and caveolin1 (p = 0.03), but positively correlated with soluble amyloid-β peptides (Aβ) concentrations (p = 0.01) and insoluble phospho-tau (S396/404, p = 0.002). No significant differences in ACE2 were observed in the 3xTgAD mouse model of tau and Aβ neuropathology. Results from immunofluorescence and Western blots showed that ACE2 protein is mainly localized in neurons in the human brain but predominantly in microvessels in the mouse brain. The present data show that an AD diagnosis is associated with higher levels of soluble ACE2 in the human brain, which might contribute to a higher risk of CNS SARS-CoV-2 infection.

## Introduction

SARS-CoV-2 (Severe Acute Respiratory Syndrome CoronaVirus 2), the cause of Coronavirus disease 2019 (COVID-19), remains a significant public health concern. Epidemiological studies have shown that fatality rates increase with age, particularly after 65 years of age [40]. In terms of death and other clinical complications, people with dementia are disproportionately impacted by COVID-19 [14, 44, 62, 66, 86]. Whether this is due to age *per se* or other factors associated with cognitive decline is currently unknown.

Although the SARS-CoV-2 virus mainly infects lower respiratory and nasopharyngeal tracts, causing respiratory failure, CNS manifestations have also been described in more than one third of hospitalized patients, especially those with severe condition [14, 46, 49, 52, 61, 85, 86]. Non-specific neurological symptoms like headache and dizziness are also commonly observed in different cohorts [33, 49] but other neurological complications reported include ischaemic stroke [8], encephalopathy [34], meningo-encephalitis [34], demyelination [87], infarcts and microhaemorrhages [26, 48].

An association between COVID-19 infection and cognitive decline due to Alzheimer’s disease (AD) or other causes is emerging. Risk factors for COVID-19 complications are often the same as those for dementia - age, obesity, cardiovascular disease, hypertension, and diabetes mellitus [44, 63, 79]. Notably, dementia *per se* is a strong predictor of COVID-19 mortality [50]. Using de-identified population-level electronic health records (EHR) from over 60 million individuals, a retrospective study showed that patients with dementia and COVID-19 had significantly worse outcomes (6-month hospitalization risk and mortality risk) than patients with dementia and no COVID-19 or patients with COVID-19 but no dementia [79]. This association remained significant after adjusting for age, sex and known COVID-19 risk factors (including, for example type 2 diabetes, cardiovascular diseases, pulmonary diseases, asthma and others). Concerns have also been raised that COVID-19 increases the risk of developing cognitive impairments, possibly secondary to cerebral ischemia in brain areas linked to cognition [3, 21, 39, 54].

Angiotensin I Converting Enzyme 2 (ACE2) is a membrane carboxypeptidase considered the main site of entry of SARS-CoV-2 into cells [38, 45, 64]. ACE2 is highly expressed in the lung, consistent with respiratory dysfunction being the first clinical consequence of the infection [15, 28]. Interestingly, ACE2 is expressed in other tissues such as the kidney, intestine, liver, testis and brain [36, 67]. The distribution of ACE2 in the brain is controversial, and original reports failed to identify the protein in the human CNS [19, 73]. Still, low levels of ACE2 mRNA were detected in the human brain using quantitative real-time RT-PCR [30]. Cerebral immunostaining was reported in endothelial and arterial smooth muscle cells [29], as well as in neurons [20]. More recently, single-cell RNA sequencing data have brought new insights on the cellular distribution of ACE2 transcripts in the brain vasculature. According to the Betsholtz mouse database, the expression of ACE2 is very high in microvascular mural cells (such as pericytes and venous vascular smooth muscle cells), but not in endothelial cells [32, 57, 77]. However, other databases report ACE2 mRNA expression in endothelial cells of mice [15, 88, 89]. So far, the available data suggest that the expression of ACE2 is lower in both endothelial cells and pericytes in the human brain compared to the mouse brain, albeit with important interregional variability [15, 31, 51, 84, 85]. In previous outbreaks of SARS-CoV, which also use ACE2 as an entry point, the virus was detected in the brains of infected patients but reported almost exclusively in neurons [18, 29, 58, 71]. Based on the above evidence, ACE2 expression in cells forming the neurovascular unit could provide a means by which SARS-CoV-2 enters the CNS.

Here, to investigate whether ACE2 levels in the brain could be associated with cognitive dysfunction, we compared mRNA and protein levels of ACE2 in *postmortem* brain samples from individuals of two different cohorts, including subjects diagnosed with AD. In the first cohort from the Religious Order Study (n=60), ACE2 protein levels were evaluated according to (i) the clinical diagnosis of no cognitive impairment (NCI), mild cognitive impairment (MCI), or AD, (ii) the neuropathological diagnosis of AD (ABC scoring) and the *antemortem* assessment of cognitive function. Associations between ACE2 and neurovascular markers were also examined. In the second cohort from other US sources (n=82), the relationships between brain ACE2 and mRNA concentrations were investigated in individuals with a Braak-based neuropathological AD diagnosis. Finally, we compared the cellular localization of ACE2 between human and mouse brains and assessed ACE2 levels in a triple transgenic mouse model of AD neuropathology, the 3xTg-AD mouse.

### Materials and methods

### Human samples

#### Cohort #1

Gray matter samples from the Brodmann area 7 (BA7) corresponding to the posterior parietal cortex were obtained from participants in the Religious Orders Study *(Rush Alzheimer’s Disease Center)*, an extensive longitudinal clinical and pathological study of aging and dementia [5, 6]. Each participant enrolled without known dementia and underwent annual structured clinical evaluations until death. A total of 21 cognitive performance tests were performed for each subject. At the time of death, a neurologist, blinded to all *postmortem* data, reviewed clinical data and rendered a summary diagnostic opinion regarding the clinical diagnosis proximate to death. Participants thus received a clinical diagnosis of NCI (*n* = 20) or MCI (*n* = 20) or AD (*n* = 20), as previously described [5–7]. The neuropathological assessment for the subjects included in the present study was performed using the ABC scoring method found in the revised National Institute of Aging – Alzheimer’s Association (NIA-AA) guidelines for the neuropathological diagnosis of AD [56]. Three different neuropathological parameters were evaluated for each subject: (A) the Thal score assessing phases of Aβ plaque accumulation [72], (B) the Braak score assessing neurofibrillary tangle pathology [13] and (C) the CERAD score assessing neuritic plaque pathology [55]. These scores were then combined to obtain an ABC score, reported as AX, BX and CX with X ranging from 0 to 3 for each parameter [56]. To determine a dichotomic neuropathological diagnosis in accordance with the revised NIA-AA guidelines [56], participants with intermediate or high levels of AD neuropathological markers were classified as AD, whereas participants with no or a low level of AD neuropathological changes were classified as Controls. Relevant data from the ROS samples used here are summarized in Table 1.

**Table 1:**
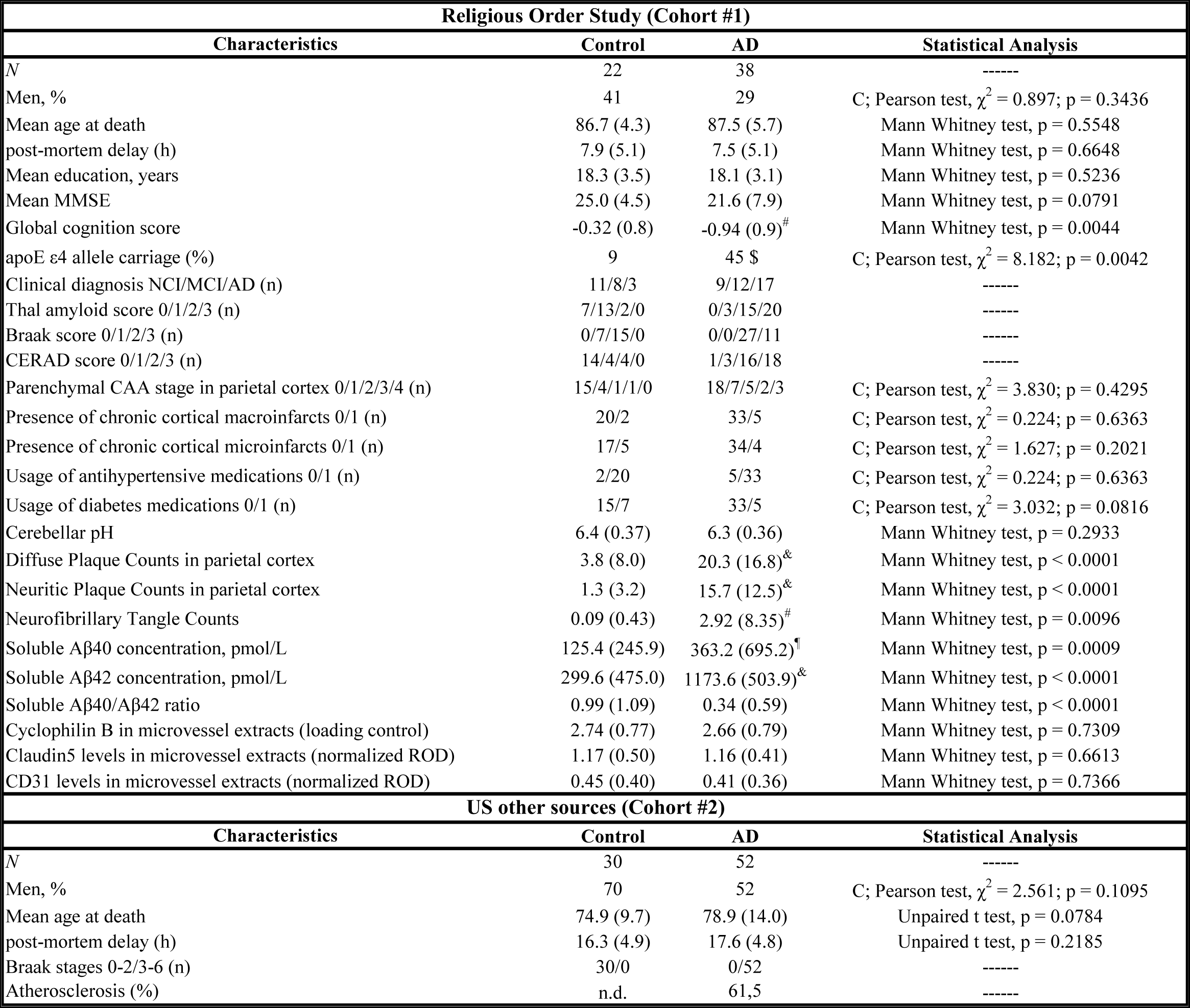
Characteristics of Cohort #1 (Religious Order Study) and Cohort #2 (US other sources). Cohort #1 characteristics (Religious Order Study): Participants were assigned to the “Control” or “AD” group based on the level of AD neuropathological changes associated with their ABC scores [70]. ABC scores were converted into one of the four levels of AD neuropathological changes (not, low, intermediate, or high) using the chart described in the revised NIA-AA guidelines [70]. Intermediate or high levels of AD neuropathological changes were assigned to the “AD” group, while those with no or a low level of AD neuropathological changes were rather assigned to the “Control” group [70]. Parenchymal CAA stages in parietal cortex were determined in the angular gyrus. Brain pH was measured in cerebellum extracts. Soluble Aβ peptide concentrations were determined by ELISA in whole homogenates of inferior parietal cortex. Values are expressed as means (SD) unless specified otherwise. Statistical analysis (compared to controls): Mann Whitney test: #p < 0.01; ¶p < 0.001; &p < 0.0001; Pearson test: £p < 0.01. Claudin5 and CD31 data in microvessel extracts were normalized with cyclophilin B as a loading control. Cohort #2 characteristics (Other US Sources): Brain samples of this cohort were provided by Harvard Brain Tissue Resource Center (Boston), Miller School of Medicine (Miami) and Human Brain and Spinal Fluid Resource Center (Los Angeles). Participants were assigned to the “Control” or “AD” group based on the Braak score. Values are expressed as means (SD). Statistical analysis (compared to controls): Unpaired t test, Pearson test. *Abbreviations: AD, Alzheimer’s disease; C, contingency; CAA, cerebral amyloid angiopathy; CERAD, Consortium to Establish a Registry for Alzheimer’s Disease; MCI, mild cognitive impairment; MMSE, Mini- Mental State Examination; NCI, healthy controls with no cognitive impairment; ROD, relative optical density*.

| Religious Order Study (Cohort #1) |  |  |  |
| --- | --- | --- | --- |
| Characteristics | Control | AD | Statistical Analysis |
| N | 22 | 38 | ----- |
| Men, % | 41 | 29 | C; Pearson test, $\chi^2 = 0.897$; p = 0.3436 |
| Mean age at death | 86.7 (4.3) | 87.5 (5.7) | Mann Whitney test, p = 0.5548 |
| post-mortem delay (h) | 7.9 (5.1) | 7.5 (5.1) | Mann Whitney test, p = 0.6648 |
| Mean education, years | 18.3 (3.5) | 18.1 (3.1) | Mann Whitney test, p = 0.5236 |
| Mean MMSE | 25.0 (4.5) | 21.6 (7.9) | Mann Whitney test, p = 0.0791 |
| Global cognition score | -0.32 (0.8) | -0.94 (0.9)<sup>#</sup> | Mann Whitney test, p = 0.0044 |
| apoE ε4 allele carriage (%) | 9 | 45 | C; Pearson test, $\chi^2 = 8.182$; p = 0.0042 |
| Clinical diagnosis NCI/MCI/AD (n) | 11/8/3 | 9/12/17 | ----- |
| Thal amyloid score 0/1/2/3 (n) | 7/13/2/0 | 0/3/15/20 | ----- |
| Braak score 0/1/2/3 (n) | 0/7/15/0 | 0/0/27/11 | ----- |
| CERAD score 0/1/2/3 (n) | 14/4/4/0 | 1/3/16/18 | ----- |
| Parenchymal CAA stage in parietal cortex 0/1/2/3/4 (n) | 15/4/1/1/0 | 18/7/5/2/3 | C; Pearson test, $\chi^2 = 3.830$; p = 0.4295 |
| Presence of chronic cortical macroinfarcts 0/1 (n) | 20/2 | 33/5 | C; Pearson test, $\chi^2 = 0.224$; p = 0.6363 |
| Presence of chronic cortical microinfarcts 0/1 (n) | 17/5 | 34/4 | C; Pearson test, $\chi^2 = 1.627$; p = 0.2021 |
| Usage of antihypertensive medications 0/1 (n) | 2/20 | 5/33 | C; Pearson test, $\chi^2 = 0.224$; p = 0.6363 |
| Usage of diabetes medications 0/1 (n) | 15/7 | 33/5 | C; Pearson test, $\chi^2 = 3.032$; p = 0.0816 |
| Cerebellar pH | 6.4 (0.37) | 6.3 (0.36) | Mann Whitney test, p = 0.2933 |
| Diffuse Plaque Counts in parietal cortex | 3.8 (8.0) | 20.3 (16.8)<sup>&</sup> | Mann Whitney test, p < 0.0001 |
| Neuritic Plaque Counts in parietal cortex | 1.3 (3.2) | 15.7 (12.5)<sup>&</sup> | Mann Whitney test, p < 0.0001 |
| Neurofibrillary Tangle Counts | 0.09 (0.43) | 2.92 (8.35)<sup>#</sup> | Mann Whitney test, p = 0.0096 |
| Soluble Aβ40 concentration, pmol/L | 125.4 (245.9) | 363.2 (695.2)<sup>¶</sup> | Mann Whitney test, p = 0.0009 |
| Soluble Aβ42 concentration, pmol/L | 299.6 (475.0) | 1173.6 (503.9)<sup>&</sup> | Mann Whitney test, p < 0.0001 |
| Soluble Aβ40/Aβ42 ratio | 0.99 (1.09) | 0.34 (0.59) | Mann Whitney test, p < 0.0001 |
| Cyclophilin B in microvessel extracts (loading control) | 2.74 (0.77) | 2.66 (0.79) | Mann Whitney test, p = 0.7309 |
| Claudin5 levels in microvessel extracts (normalized ROD) | 1.17 (0.50) | 1.16 (0.41) | Mann Whitney test, p = 0.6613 |
| CD31 levels in microvessel extracts (normalized ROD) | 0.45 (0.40) | 0.41 (0.36) | Mann Whitney test, p = 0.7366 |
| US other sources (Cohort #2) |  |  |  |
| Characteristics | Control | AD | Statistical Analysis |
| N | 30 | 52 | ----- |
| Men, % | 70 | 52 | C; Pearson test, $\chi^2 = 2.561$; p = 0.1095 |
| Mean age at death | 74.9 (9.7) | 78.9 (14.0) | Unpaired t test, p = 0.0784 |
| post-mortem delay (h) | 16.3 (4.9) | 17.6 (4.8) | Unpaired t test, p = 0.2185 |
| Braak stages 0-2/3-6 (n) | 30/0 | 0/52 | ----- |
| Atherosclerosis (%) | n.d. | 61,5 | ----- |

#### Cohort #2

Gray matter samples from the parietal cortex were obtained from 3 different institutions in the United States: 1- *Harvard Brain Tissue Resource Center,* Boston, Massachussetts, 2- *Brain Endowment Bank*, Miami, Florida. 3- *Human Brain and Spinal Fluid Resource Center*, Los Angeles, California [23]. All 82 parietal cortex samples were from the Brodmann area 39 (BA39), corresponding to the inferior region of the parietal cortex. Neuropathological diagnoses were based on Braak scores that were available for all cases. Braak scores of I or II were classified as Controls while Braak scores of III, IV, V and VI were considered as AD (Table 1).

### Immunostaining

To demonstrate ACE2 localization in *postmortem* human brain tissue, we tested a number of commercially available antibodies using a wide range of immunostaining protocols. Immunostaining was performed on formalin-fixed, paraffin-embedded (FFPE) tissue sections (6 µm) of human parietal cortex, on fresh-frozen (FF) tissue sections (12 µm) from human and mouse hippocampus, as well as on human and murine isolated brain microvessels (see below). Briefly, FFPE sections were deparaffinized in CitriSolv hybrid and rehydrated with decreasing concentrations of ethanol in water. Antigen retrieval was then performed by boiling slides in Tris buffer (10 mM, pH 9.0) with 1 mM EDTA and 0.05% (v/v) Tween-20 in a microwave for 15 minutes and letting them cool for 30 minutes at room temperature. Sections were quenched with 50 mM NH_4_Cl, digested with trypsin 0.1% (w/v, Sigma-Aldrich) at 37°C for 15 min, and incubated in Tris-buffered saline (TBS) with 0.3 M glycine for 15 minutes. Sections were then blocked sequentially with Bloxall, avidin/biotin blocking kit (Vector Laboratories, CA) and Superblock (Thermo) with 0.2% Triton-X100, and used for immunohistochemistry. FF sections and brain microvascular fractions were kept at -80°C until use, then vacuum-dried at 4°C and fixed in 4% (w/v) paraformaldehyde (pH 7.4) for 20 min at room temperature. All sections were then blocked and permeabilized for 1h with Superblock containing 0.2% (v/v) Triton X-100. Incubation with primary antibodies (various rabbit anti-ACE2, mouse anti-NeuN MAB377, goat anti-collagen IV AB789) was performed overnight at 4°C in Superblock with 0.05% Tween-20. For immunohistochemistry, after multiple washing in PBS, sections were incubated with biotinylated secondary antibodies (Jackson Immunoresearch) and then with streptavidin-HRP (ABC Elite kit). ACE2 localization was revealed using the ImmPACT AMEC red substrate and nuclei were counterstained with Mayer’s hematoxylin. For immunofluorescence, after washes, secondary antibodies (conjugated to Alexa Fluor 555, 647 and 750, which use channels with less autofluorescence) were added to sections for 1h. Slides were then sequentially incubated with 4′,6- diamidino-2-phenylindole (DAPI) and TrueBlack Plus (Biotium, CA). Photomicrographs were recorded with a Cytation 5 or EVOS fl Auto Imaging System (Thermo Fisher).

### Protein fractionation from human parietal cortex homogenates

Each inferior parietal cortex sample (∼100 mg) from the Cohort #1 was sequentially sonicated and centrifuged to generate two protein fractions: a Tris-buffered saline (TBS)-soluble fraction containing soluble intracellular, nuclear and extracellular proteins and a detergent-soluble protein fraction containing membrane-bound proteins extracted with a mix of detergents (0.5% Sodium dodecyl sulfate (SDS), 0.5% deoxycholate, 1% Triton), as previously reported [74, 75]. For samples of the Cohort #2, frozen extracts from the parietal cortex were ground into a fine powder on dry ice with a mortar and pestle. Approximatively 50 mg of this fine powder was used for total protein extraction. Then, a lysis buffer (50 mM Tris-HCL to pH 7.4, 150 mM NaCl, 1% triton and 0.5% sodium deoxycholate) containing protease (complete 25 X and Pepstatin A) and phosphatase inhibitors (sodium fluoride and sodium vanadate) was added in the proportion of 4 times the sample weight (400 μL for 50 mg). The sample solution was homogenized on ice by sonication using a Sonic Dismembrator (Fisher, Pittsburgh, PA) with two 10-second pulses and a 30-second stop between steps. Samples were centrifuged 20 minutes at 10,000 g at 4 °C. Protein contents of supernatants were quantified using a bicinchoninic acid assay (Thermofisher cat: P123227). Protein homogenates in Laemmli were prepared as described below.

### Isolation of human brain microvessels

The method used to generate microvessel-enriched extracts from frozen human parietal cortex samples has been described in our previous publications [10–12]. Briefly, this method consists of a series of centrifugation steps, including one density gradient centrifugation with dextran, after which the tissue is filtered through a 20-µm nylon filter. Two fractions were generated: one enriched in cerebral microvessels (isolated microvessel-enriched fraction) and the other consisting of microvessel-depleted parenchymal cell populations. Cerebral fractions enriched and depleted in endothelial cells were evaluated using immunoblotting of vascular and neuronal markers, as shown previously [11]. Isolation of brain microvessels was performed on human samples of Cohort #1 (final n = 57) and the fractions were used for immunostaining and Western blot analysis.

### RNA extraction and RT-qPCR analysis

As mentioned above, powderized parietal cortex tissues of Cohort #2 were kept at -80°C. Approximately 100 mg of this fine powder was used for total RNA extraction with TRIzol (Ambion). Unless otherwise noted, all steps were performed on ice or at 4°C. Samples were homogenized by sonication with a Sonic Dismembrator (Fisher, Pittsburgh, PA) with 4-sec pulses in 500 μL of TRIzol. Chloroform (100 μL) was added to the solution, mixed and incubated for 2 minutes before centrifugation at 12,000 g for 15 minutes. Supernatants were kept, 250 μL of isopropanol were added followed by a 10-min incubation at -80°C and a 10-min centrifugation, at 12,000 g. The pellet was resuspended in 500 μL of ethanol 75 % and centrifuged 5 minutes at 7500 g. The dried pellet was resuspended in 80 μL RNAse-free water and incubated 10 minutes at 57 °C. The RNA concentration was measured with an Infinite F200 (Tecan). The reverse transcription (RT) was performed with 1 µg of RNA. As a first step, genomic DNA was removed following the AccuRT Genomic DNA removal protocol (Applied Biological Materials, ABM, Vancouver, Canada). Then the RT master mix (ABM) was added to RNA samples and incubated (10 min at 25°C, 50 min at 42°C and 5 min at 85°C) as per the manufacturer’s protocol. All qPCR experiments were performed on the LightCycler 480 (Roche) with the BrightGreen mix (ABM) and primers at 10 µM. After enzyme activation for 10 min at 95°C, 50 cycles were performed (15 sec at 95°C and 1 min at 60°C), followed by 1 sec at 95°C and 1 min at 45°C. Reference gene GAPDH (primers Forward: TCTCCTCTGACTTCAACAGCGAC and Reverse

:CCCTGTTGCTGTAGCCAAATTC) was used to normalize the mRNA expression. The relative amounts of each transcript were calculated using the comparative Ct (2-ΔΔCt) method. For *Ace2* qPCR (primers Forward: GTGCACAAAGGTGACAATGG and Reverse: GGCTGCAGAAAGTGACATGA), 12 controls and 19 AD individuals were used. We used a cut- off of ≥ 35 cycles for these samples.

### Isolation of murine brain microvessels and protein fractionation

Four (4) or six (6)-, 12- and 18-month-old 3xTg-AD (APPswe, PS1M146V, tauP301L) mice produced at our animal facility were used in equal numbers of males and females in each group. These mice show progressive accumulation of Aβ plaques and neurofibrillary tangles which are detectable at 12 months and are widespread after 18 months [9, 16]. Mice from our colony were fed a standard chow (Teklad 2018, Harlan Laboratories, Canada) from breeding to 5 months of age. For microvessels extraction, mice were then fed a control diet (CD; 20%kcal from fat) or a high-fat diet (HFD; 60%kcal from fat) from 6 to 18 months of age, in order to worsen neuropathology, memory performance and also induce metabolic impairments [4, 41, 68, 76], which are associated with a higher risk of developing severe SARS-CoV-2 infections [1, 40].

The protein extraction method results in a TBS-soluble fraction (intracellular and extracellular fraction), a detergent-soluble fraction (membrane fraction) as previously described [70].

Brain microvessels from 3xTg-AD mice were generated with a protocol similar to the one used for frozen human brain samples, as reported previously [10–12] (see Supplementary material). All experiments were performed in accordance with the Canadian Council on Animal Care and were approved by the Institutional Committee at the Centre Hospitalier de l’Université Laval (CHUL).

### Western blot analysis

Protein homogenates from human parietal cortex and murine whole brain extracts were added to Laemmli’s loading buffer and heated 10 min at 70°C. TBS- and detergent-soluble fractions from homogenates of human parietal cortex were also added to Laemmli’s loading buffer and heated 5 min at 95°C. Equal amounts of proteins per sample (8 µg for both human and murine brain microvessel extracts and 12 µg for protein homogenates of human parietal cortex, 15 µg for protein homogenates of mouse brain) were resolved by sodium dodecyl sulphate-polyacrylamide gel electrophoresis (SDS-PAGE). All samples, loaded in a random order, were run on the same immunoblot experiment for quantification. Proteins were electroblotted on PVDF membranes, which were then blocked during 1h with a PBS solution containing 5% non-fat dry milk, 0.5% BSA and 0.1% Tween-20. Membranes were then incubated overnight at 4°C with primary antibodies (rabbit anti-ACE2, #ab108252, 1:1000, rabbit anti-TMPRSS2 #ab109131, 1:1000).

Membranes were then washed three times with PBS containing 0.1% Tween-20 and incubated during 1h at room temperature with the secondary antibody (goat/donkey anti-rabbit HRP Jackson ImmunoResearch Laboratories, West Grove, PA; 1:60,000 or 1:10,000 in PBS containing 0.1% Tween-20 and 1% BSA). Densitometric analysis was performed using ImageLab (Bio-Rad). Uncropped gels of human samples immunoblot assays are shown in the Supplementary Material (Figure S5 and Figure S6).

### Data and statistical analysis

An unpaired Student’s t-test was performed when only two groups were compared, with a Welch correction when variances were not equal. If the data distribution of either one or both groups failed to pass the normality tests (Shapiro-Wilk test or Kolmogorov-Smirov test), groups were compared using a non-parametric Mann-Whitney test. When more than two groups were compared, parametric one-way ANOVA followed by Tukey’s multiple comparison tests or two-way ANOVA were used. If criteria for variance (Bartlett’s) or normality were not met, non-parametric Kruskal-Wallis ANOVA followed by Dunn’s multiple comparison tests were used. If needed, data were log transformed to normalize distributions. For all data, statistical significance was set at P < 0.05. Individual data were excluded for technical reasons or if determined as an outlier using the ROUT (1%) method in GraphPad Prism. Statistical analysis was done by Pearson correlation to analyze the correlation between ACE2, *antemortem* evaluation and different proteins. All statistical analyses were performed with Prism 9 (GraphPad, San Diego, CA, USA) or JMP (version 16; SAS Institute Inc., Cary, IL) software.

## Results

### Association between ACE2 in the parietal cortex, the neuropathological diagnosis of AD and cognitive scores

Table 1 summarizes the clinical and biochemical data from Cohorts #1 (Religious Order Study, ROS) and #2 (Other US sources). ACE2 proteins from parietal cortex samples of each subject were fractionated into TBS-soluble (cytosolic, extracellular, nuclear and secreted proteins), detergent-soluble (membrane-bound proteins) and microvessel-enriched fractions (vascular proteins). Representative Western immunoblots of ACE2 and analyses are shown in Figure 1 for Cohort #1 and Figure 2 for Cohort #2. A band migrating at approximately 100 kDa, corresponding to full-length ACE2, was observed in each fraction (Figure 1A,E,I, Figure 2A).

**Figure 1:**
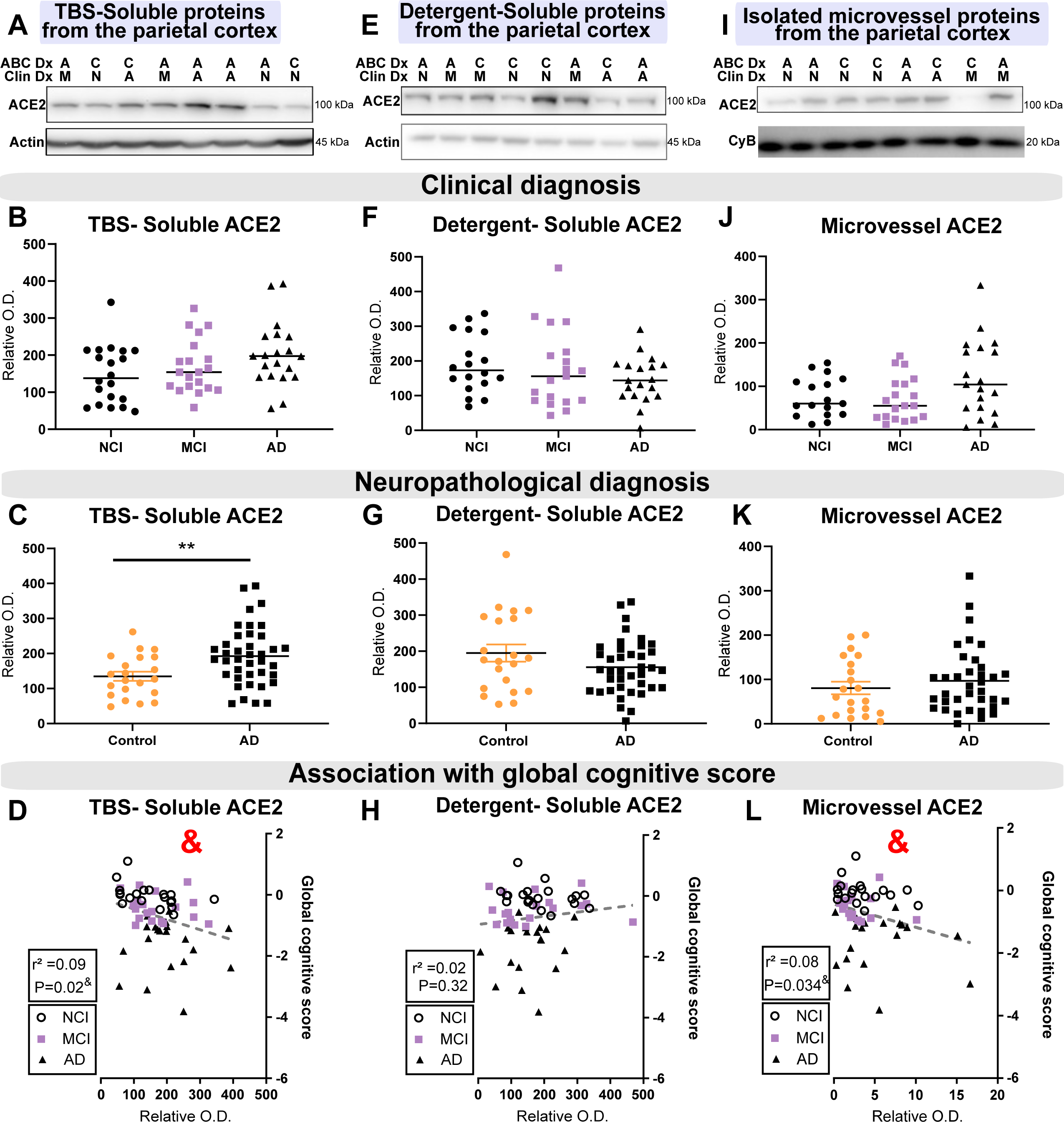
Levels of TBS-Soluble ACE2 protein are higher in AD individuals and are negatively correlated with global cognitive score. Parietal cortex levels of ACE2 protein from Cohort #1 were determined by Western Blot in three fractions: a TBS-soluble fraction (A-D), a detergent-soluble fraction (E-H) and a microvessel-enriched fraction (I-L). No statistical difference was detected for ACE2 in the three fractions when subjects were classified according to *antemortem* clinical diagnosis (B, F, J). Levels of the ACE2 protein were higher in the TBS-soluble fraction in individuals with a neuropathological diagnosis of AD based on ABC scoring (C, G, K). TBS-soluble ACE2 and microvascular ACE2 levels were negatively correlated with the global cognitive score (D, L). An equal amount (12 µg) of proteins per sample for both TBS-soluble and detergent soluble fractions was loaded and 8 µg of proteins per sample was loaded for microvessel-enriched fractions. All samples, loaded in a random order, were run on the same gel and transferred on the same membrane before immunoblotting for quantification. Examples were taken from the same experiment, and consecutive bands loaded in random order are shown. Actin and cyclophilin B are shown as loading controls. Data are represented as a scatterplot. Horizontal lines indicate mean ± SEM. Statistical analysis: Mann-Whitney test **p < 0.01, Coefficient of determination &p < 0.05. *Abbreviations: ACE2, Angiotensin-Converting Enzyme 2; A/AD, Alzheimer’s disease; C, control; Clin Dx, clinical diagnosis; ABC Dx, ABC neuropathological diagnosis; CyB, cyclophilin B; M/MCI, mild cognitive impairment; N/NCI, healthy controls with no cognitive impairment; O.D., optical density; TBS, Tris-Buffered Saline*.

**Figure 2:**
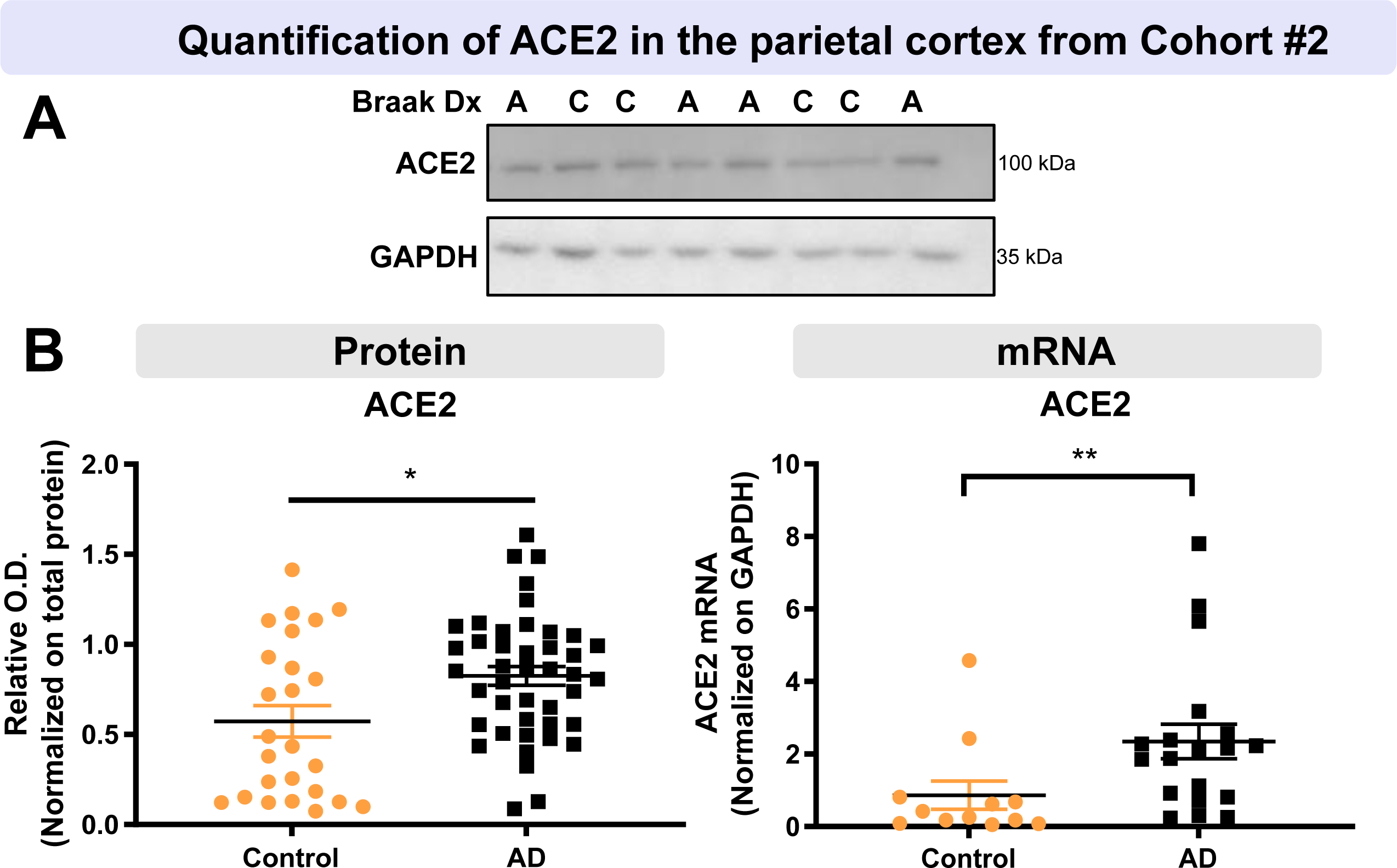
Higher ACE2 protein and mRNA levels in parietal cortex of AD participants from the second cohort. AD subjects from Cohort #2 had higher levels of ACE2 protein and mRNA compared to controls. Diagnosis was determined using Braak staging. ACE2 levels were determined by Western-Blot and qPCR analysis (A, B). Statistical analysis: Mann-Whitney test *p < 0.05, Unpaired t-test **p < 0.01. All samples, loaded in a random order, were run on the same immunoblot experiment for quantification. Examples were taken from the same immunoblot experiment, and consecutive bands loaded in random order are shown. GAPDH is shown as a loading control. Data are represented as a scatterplot. Horizontal lines indicate mean ± SEM. *Abbreviations: A, Alzheimer’s disease (Braak scores III-VI); ACE2, Angiotensin-Converting Enzyme 2; C, control (Braak scores I or II); Dx, diagnostic; O.D., optical density*.

We first evaluated whether ACE2 protein levels in extracts from the parietal cortex from 60 individuals from the Cohort #1, ROS. When the subjects were classified according to the neuropathological ABC diagnosis, higher levels of ACE2 protein were found in TBS-soluble fractions from AD subjects compared to non-AD participants (p = 0.0087) (Figure 1C). When the subjects were classified according to the clinical diagnosis, only a non-significant trend towards higher ACE2 concentrations was observed in the TBS-soluble fraction, using a non-parametric Kruskal-Wallis ANOVA (p = 0.1471) (Figure 1B). However, the difference between AD and Controls was statistically significant when this comparison was performed only in individuals with parenchymal cerebral amyloid angiopathy (p = 0.0022) (pCAA; Figure S1). On the other hand, ACE2 levels assessed in the detergent-soluble fraction, enriched for membrane-associated proteins, remained similar between groups (Figure 1E-G). We next measured ACE2 protein levels in microvessel extracts. Given the high interindividual variability, only a non-significant trend towards increased ACE2 protein levels was observed in individuals with an AD clinical diagnosis (p = 0.1712) (Figure 1J, -K).

Interestingly, the levels of ACE2 found in TBS-soluble and microvessel-enriched fractions were inversely associated with *antemortem* global cognitive scores (Figure 1D, L and Figure 3). This association remained significant after adjustment for age at death and sex.

**Figure 3:**
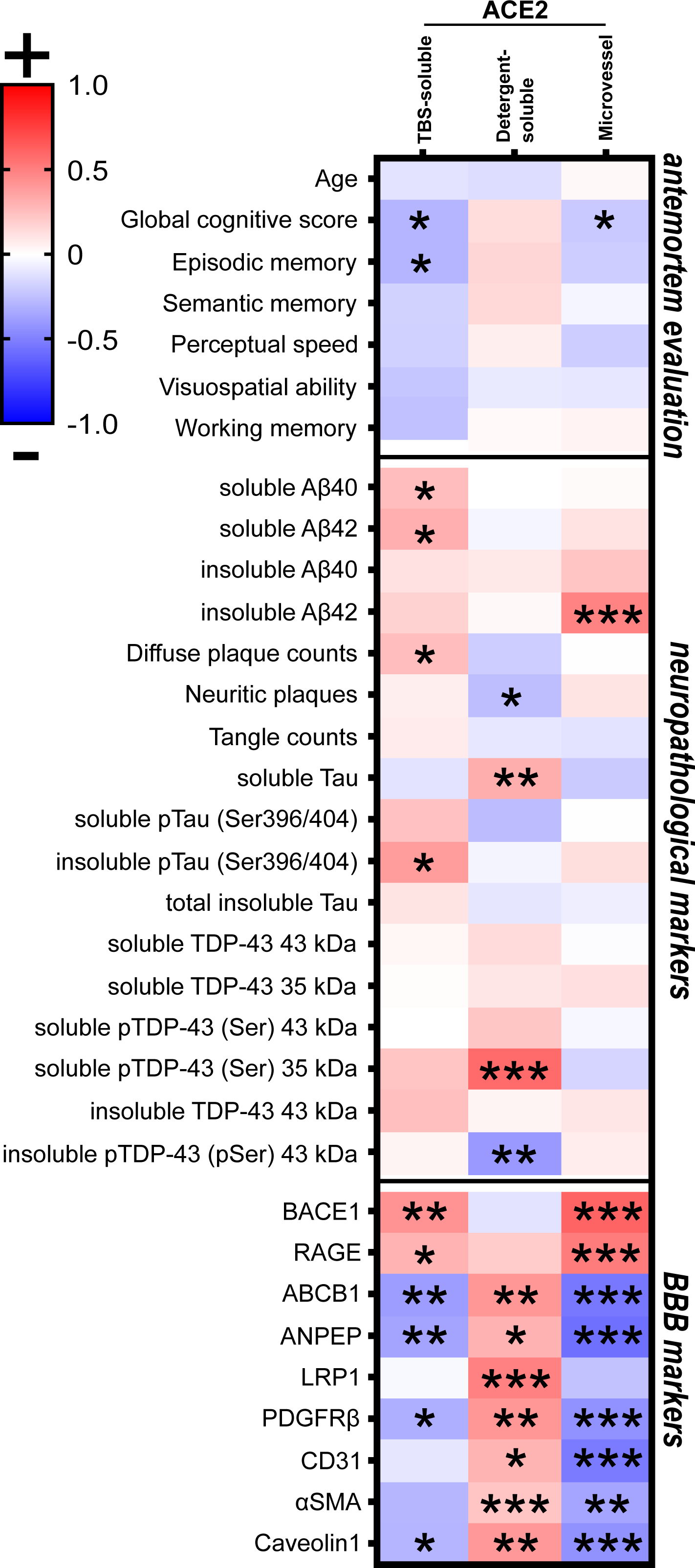
ACE2 protein in TBS-soluble/microvascular protein fractions show opposite relationships with detergent-soluble ACE2 when correlating with AD markers. Heat-map of hierarchical clustering analysis of correlation coefficients from partial correlation analyses with *antemortem* evaluation, neuropathological markers (soluble and insoluble proteins of the parietal cortex) and BBB markers (microvascular fractions of the parietal cortex). The significance of the correlation (inverse in blue, positive in red) between two elements is entered in the associated box. Statistical analysis: Coefficient of determination *p < 0.05, **p < 0.01 and ***p < 0.001. All proteins presented in these heat-maps were determined by Western-blot analysis except Aβ peptides which were quantified using ELISA. Total soluble and insoluble Tau was detected using Tau (640-680) antibody. AD2 antibody recognized Tau phosphorylated at S396. Phosphorylated TDP43 antibody recognized at pSer409/410. *Abbreviations: ABCB1, ATP Binding Cassette Subfamily B Member 1; ACE2, Angiotensin-Converting Enzyme 2; ANPEP, Aminopeptidase N; BBB, blood-brain barrier; BACE1, Beta-Secretase 1; CD31 or PECAM1, Platelet endothelial cell adhesion molecule; LRP1, Low density lipoprotein receptor-related protein 1; NS, non significative; PDGFRß, Platelet Derived Growth Factor Receptor Beta; RAGE, Receptor for Advanced Glycation Endproducts; αSMA, alpha smooth muscle actin; TDP-43, TAR DNA binding protein 43; TBS, Tris-Buffered Saline*.

To corroborate these results, we performed Western immunoblots on a second series of human brain samples from the Cohort #2 [23] (Figure 2A). Consistently, higher levels of ACE2 protein were detected in individuals with a neuropathological diagnosis of AD (Figure 2B). In addition, *Ace2* mRNA levels were significantly higher in individuals with a Braak-based diagnosis of AD compared to Controls (Figure 2B), suggesting a regulation at the transcriptional level. No difference was observed in the levels of transmembrane protease serine type 2 (TMRPSS2) (Figure S2), a protein that plays a key role in SARS-CoV-2 infection by activating the spike protein, facilitating entry into target cells using ACE2 [38].

### TBS-soluble ACE2 is positively associated with clinical, neuropathological, and vascular markers of AD, while detergent-soluble ACE2 displays opposite trends

Hierarchical clustering of correlation coefficients (strength of association) was performed to identify variables associated with differences in ACE2 in Cohort #1. Although the leading risk factor for AD is age, no significant correlation was found between ACE2 levels in all fractions tested and the ages of death (Figure 3, Figure S3), which were equivalent between groups (Table 1). These observations suggest that the greater soluble ACE2 in individuals with AD in the ROS cohort was not driven by age. However, the age interval (74-98 years) was too small to detect an effect of aging *per se* on cerebral ACE2. Beside the inverse association with global *antemortem* cognitive scores of participants (r^2^ = -0.09, P < 0.05, Figure 1D and Figure 3), higher *postmortem* TBS-soluble concentrations of ACE2 were also significantly correlated with failing episodic memory, a domain predominantly affected in AD (Figure 3).

Associations were then examined with neuropathological markers of AD, previously assessed in the parietal cortex from the sample series (Figure 3). TBS-Soluble ACE2 levels were positively associated with markers that are greater in AD like diffuse plaque counts, soluble Aβ levels and insoluble phospho-tau (pS396/404 epitope) (Figure 3). In contrast, levels of detergent-soluble ACE2 were negatively associated with the insoluble phosphorylated form of TDP-43 (which is higher in AD [12]) but positively with soluble phospho-TDP-43 C-terminal fragment migrating at approximately ∼35 kDa (which is lower in AD [12]) and soluble tau (Figure 3). Similarly, neurovascular proteins such as platelet-derived growth factor receptor β (PDGRFRβ), ABCB1 and Caveolin1 correlated positively with membrane-bound ACE2 but inversely with TBS-soluble and vascular forms of ACE2, which in turn correlate with ß-secretase 1 (BACE1) and Advanced glycosylation end product-specific receptor (RAGE), proteins involved in the formation and accumulation of Aβ (Figure 3).

Single-cell RNA sequencing data in the mouse and human brain show that ACE2 mRNA expression is enriched in pericytes [31, 57], and PDGFRβ, a pericyte marker, is reduced in AD [12, 53]. Here, we found that microvascular PDGFRβ and Aminopeptidase N (ANPEP) levels were negatively correlated with TBS-soluble ACE2 levels but were positively associated with detergent- soluble ACE2 levels (Figure 3, Figure S3), suggesting a possible release of ACE2 from membranes linked with pericyte-related dysfunctions at the blood-brain barrier (BBB).

Together, these results suggest that the elevation of ACE2 in TBS-soluble and, to a lesser extent, in microvessel fractions are associated with more advanced Aβ and tau pathologies and with a pattern of changes in vascular proteins consistent with AD progression. By contrast, membrane-bound ACE2 exhibited opposite trends and was strongly associated with reduced TDP-43 proteinopathy and consolidated BBB markers.

### ACE2 is observed in human and murine neurons and cerebral vessels

Localizing ACE2 within the neurovascular unit at the interface between the blood and the brain provides basic information about SARS-CoV-2 penetration into the CNS. Therefore, we sought to determine whether ACE2 protein was enriched in brain microvessel extracts compared to post-vascular parenchymal fractions and unfractionated homogenates from human (parietal cortex) and mouse (whole brain) samples (Figure 4). We found a strong enrichment of ACE2 in murine microvessels, along with endothelial (Claudin5) and pericyte (PDGFRβ) markers (Figure 4F). However, in human brain samples, ACE2 protein levels were more comparable between microvessel and parenchymal fractions, the latter being enriched in the neuronal marker (synaptophysin) (Figure 4A). Thus, these Western blot results suggest that the localization of ACE2 in the brain differs between both species, with a cerebrovascular predominance in the mouse not observed in humans.

**Figure 4:**
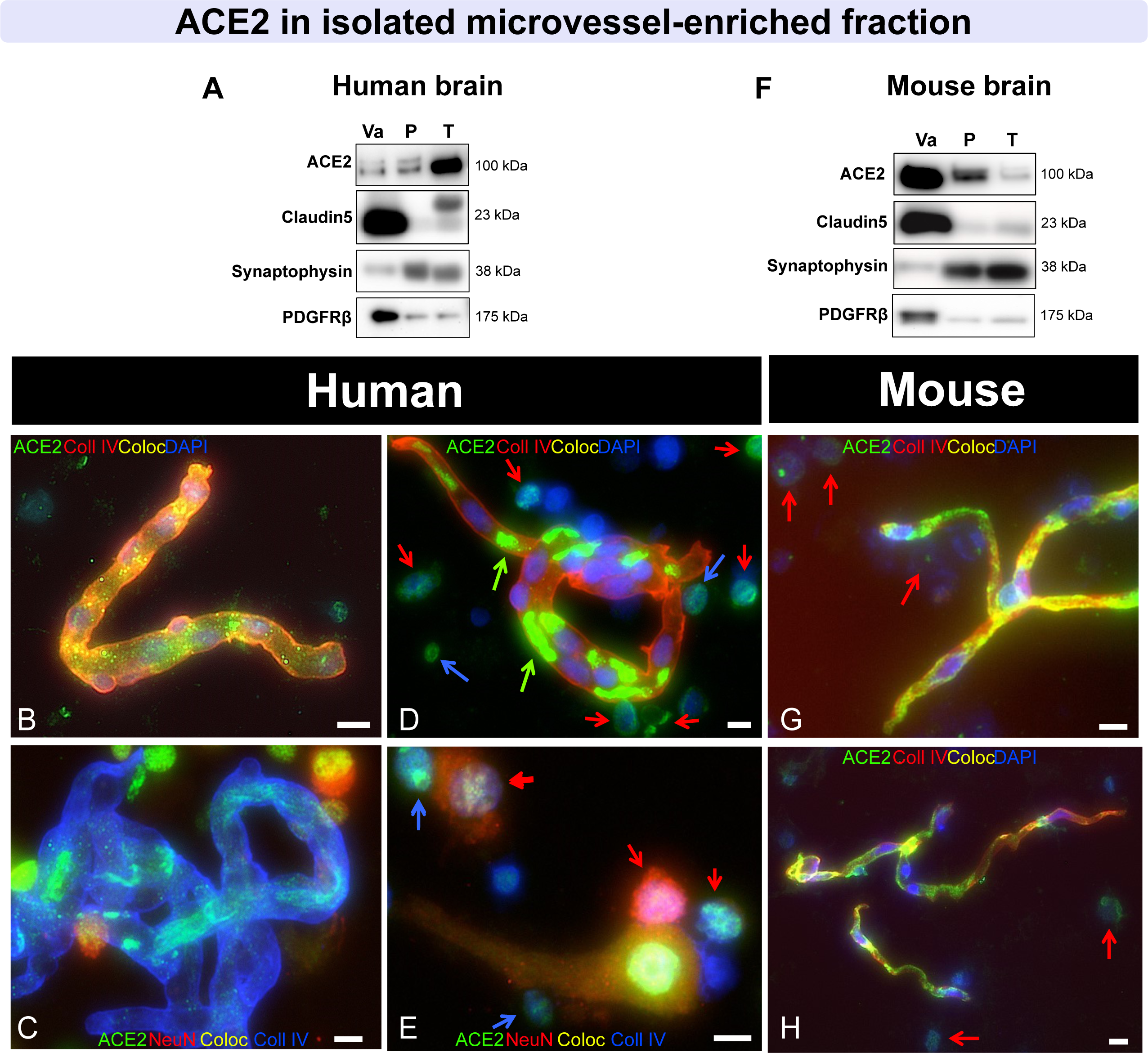
In fractionated brain homogenates, ACE2 immunosignal is predominantly observed in neurons in human samples and in the vasculature in mice. (A, F) Immunoblotting detection of ACE2 in human (parietal cortex) (A) and mouse (whole brain)(F) vascular fractions “*Va*”, compared to postvascular parenchymal samples depleted in vascular cells “*P*” and total unfractionated homogenates “*T*”. For comparison purposes, synaptophysin (a synaptic/neuronal marker), Claudin5 (endothelial marker), and PDGFRß (pericyte marker), are also shown. In the mouse brain, ACE2 is highly enriched in microvessels compared to the postvascular fraction, different to what is observed in the human (A, F). (B-E, G, H) Representative immunostaining of ACE2 (green) in human (B-E) and murine cerebrovascular fractions (G, H), with collagen IV (endothelial marker) in red (B, D, G, H) or blue (C, E), as well as NeuN (neuronal marker) in red (C, E) and DAPI (nuclei) in blue (B-H). In human samples, moderate ACE2 staining is observed in neurons, whereas vascular ACE2 staining is strong in mice. Red arrows point to ACE2+/NeuN+ cells, blue arrows to ACE2+/NeuN- cells, and green arrows to erythrocytes. ACE2 antibodies: rb mAb #ab108252 (A, F), rb pAb #HPA000288 (B, D, G) and #35-1875 (C, E, H). Scale bar: 10 µm.| *Abbreviations: ACE2, Angiotensin-Converting Enzyme 2; Coll IV, Collagen IV; Coloc, Colocalization; PDGFRß, Platelet Derived Growth Factor Receptor Beta. mAb, monoclonal antibody; pAb, polyclonal antibody*.

To confirm the cellular localization of ACE2, immunostaining was also performed on fractionated brain homogenates (Figure 4B-E, G, H). A moderate immunofluorescent signal was detected inside microvessels isolated from human brains (collagen IV-positive, Figure 4B, C) and NeuN-positive neurons (Figure 4C, E). By contrast in the mouse, ACE2 immunosignal was intense in microvessels and colocalized well with collagen IV (Figure 4G, H). To validate immunostaining in human tissue sections, nine anti-ACE2 antibodies were used in human testis samples where ACE2 is highly expressed in Leydig and Sertoli cells (Figure S4). All antibodies showed a clear signal in this tissue, but detection of ACE2 in the human brain, where levels are at least 20 times lower (results not shown), was challenging. In human hippocampal sections, ACE2 was detected in NeuN-positive neurons, particularly in large ones staining weakly for DAPI and in small ones with a strong DAPI signal (Figure 5A, B). In sections of human parietal cortex, ACE2 detection was also more prominent in neuron-like cells (Figure 5C, D, E). On the other hand, in mouse sections, ACE2 staining was more intense in the cerebrovasculature and colocalized neatly with PDGFRß, indicating an expression in mouse pericytes (Figure 5F-H). These results are consistent with Western blot data, showing that ACE2 can be detected in neuron-like cells in the human brain, whereas it is concentrated in cerebrovascular cells in the mouse.

**Figure 5:**
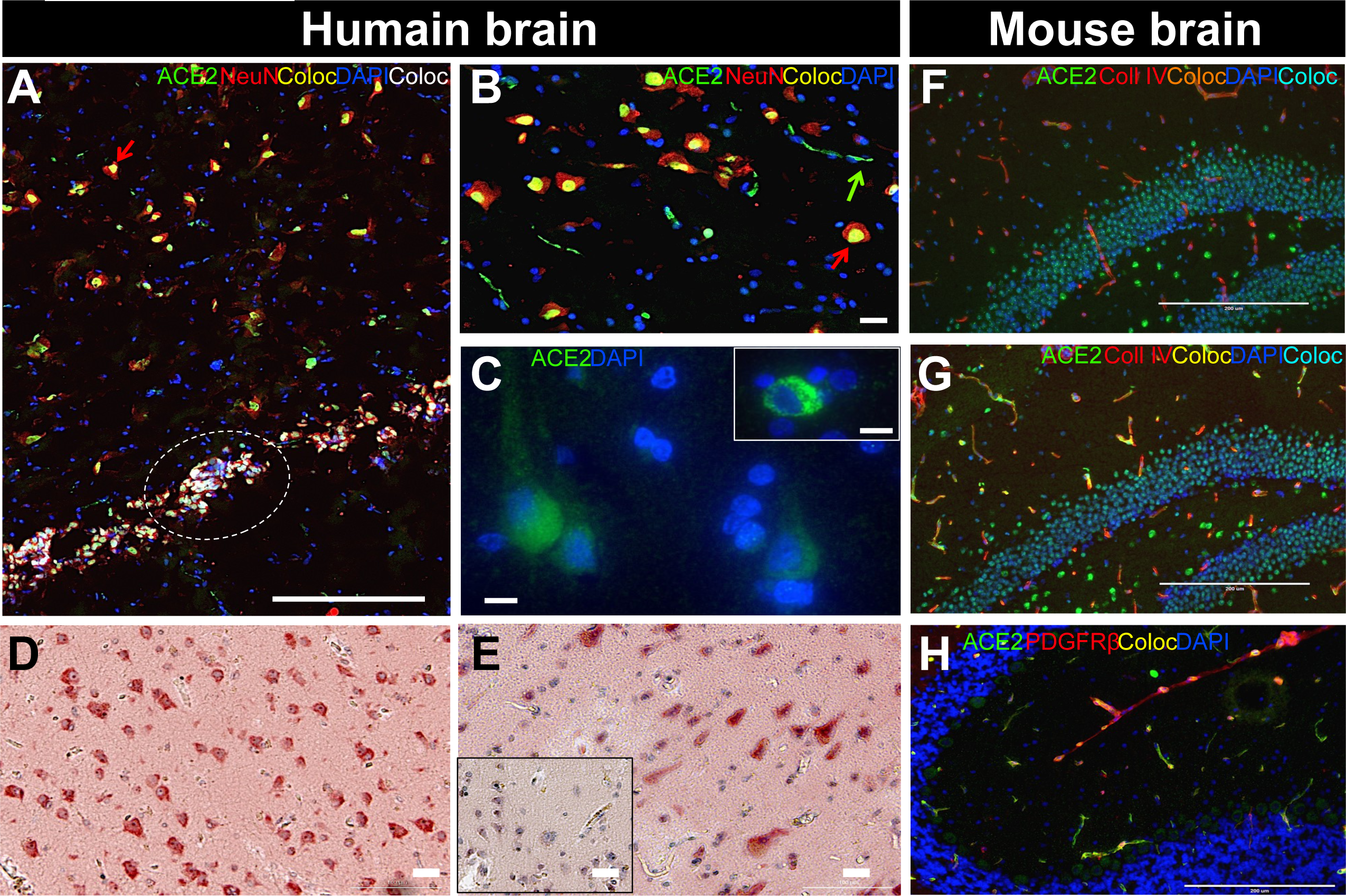
In tissue sections, ACE2 immunostaining is predominantly observed in neurons in the human brain and in the cerebrovasculature in mice. (A-E) Representative immunostaining of ACE2 (green or red) in fresh frozen human hippocampus (A, B), formalin-fixed paraffin- embedded parietal cortex (C-E), and in murine fresh frozen hippocampal and cerebellar sections (F-H). For immunofluorescence, NeuN (neuronal marker)(A, B), collagen IV (endothelial marker)(F, G) or PDGFRß (pericyte marker) (H) are in red, and DAPI (nuclei) is in blue (A-C, F- H). For immunohistochemistry, sections were counterstained with hematoxylin (nuclei)(D, E), and negative control is inserted in (E). In human samples, strong ACE2 staining is observed in large and small neurons, whereas in mice, neuronal signal is moderate and vascular ACE2 staining is strong and colocalizes well with PDGFRß. Red arrows point to ACE2+/NeuN+ cells and green arrows to erythrocytes; dashed white line highlights small ACE2+/NeuN+ cells that stain strongly for DAPI. ACE2 antibodies: rabbit mAb #ab108252 (C, G), rabbit pAb #HPA000288 (A, B, E, F) and #35-1875 (D) or goat pAb #AF3437. Scale bars: 200 µm (A, F, G, H), 20 µm (B, D, E) and 10 µm (C). *Abbreviations: ACE2, Angiotensin-Converting Enzyme 2; Coll IV, Collagen IV; Coloc, Colocalization; PDGFRß, Platelet Derived Growth Factor Receptor Beta; mAb, monoclonal antibody; pAb, polyclonal antibody*.

### Microvascular and whole-brain ACE2 protein levels are not altered in a mouse model of AD

To probe whether changes in ACE2 could be a consequence of classical tau and Aβ neuropathology, we used the triple transgenic mouse model of AD (3xTg-AD) [59], which develops Aβ plaques and neurofibrillary tangles by 12 months of age. We quantified ACE2 protein levels in both 3xTg-AD and non-transgenic mice from two different cohorts: (i) mice of 4 or 6, 12, and 18 months of age, (ii) and 18-month-old mice fed either a control or a HFD that exacerbates neuropathology [76] (Figure 6). No significant change was observed in protein levels of ACE2 in TBS-soluble or detergent-soluble fractions according to genotype and age (Figure 6A). Similarly, ACE2 in cerebrovascular fractions did not vary according to genotype, age, and diet (Figure 6B). These results suggest that the development of human tau and Aβ neuropathology in mice is insufficient to increase murine ACE2 levels, even when combined with aging and HFD, two risk factors for AD and COVID-19 infection.

**Figure 6:**
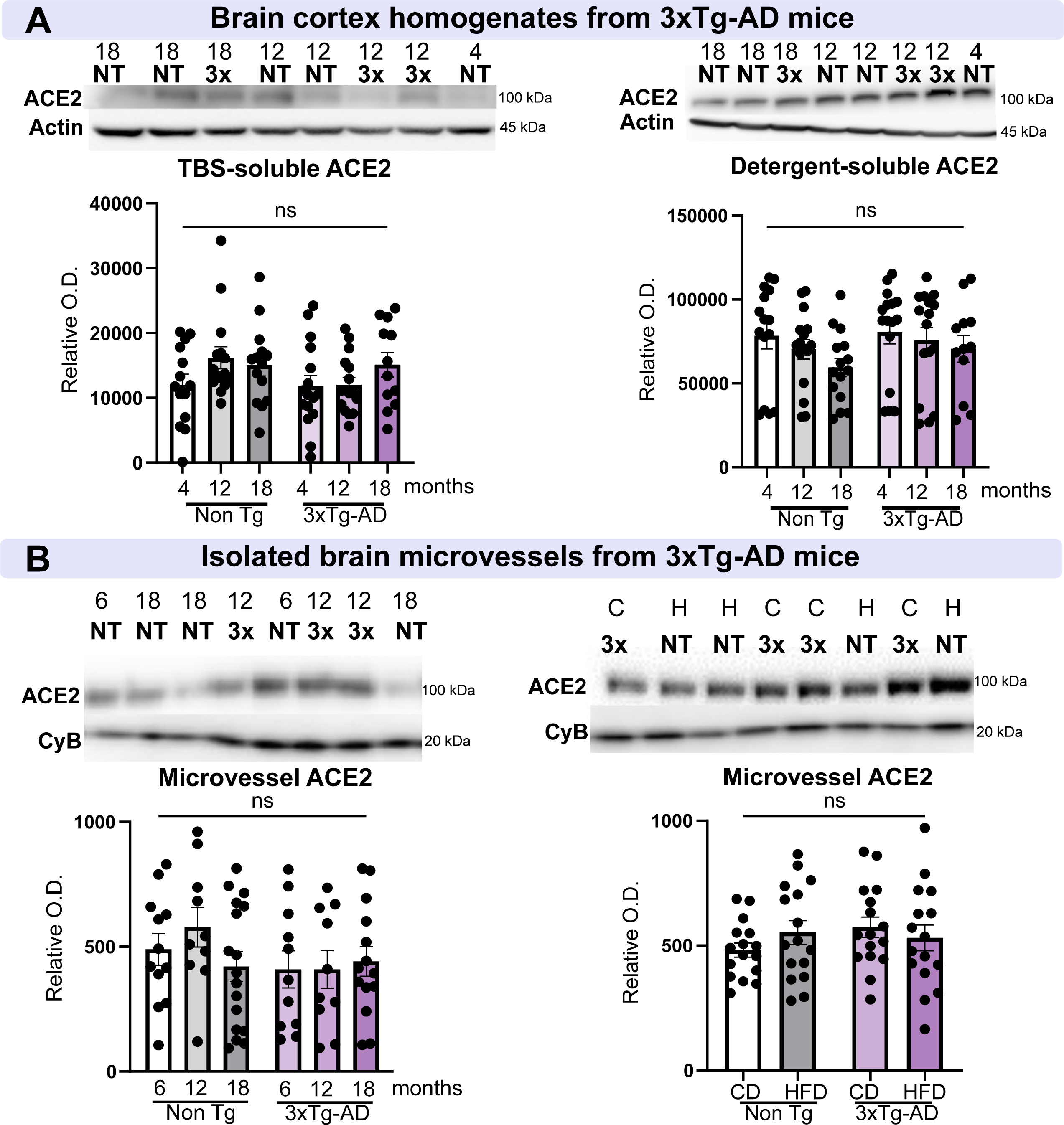
ACE2 levels are not altered in a model of AD, the 3xTg-AD mouse. (A) Determination of ACE2 levels by Western immunoblotting in brain homogenates from NonTg and 3xTg-AD mice aged 4, 12, and 18 months. No difference was observed in TBS-soluble and detergent-soluble ACE2. (B) In brain microvessel-enriched fractions from NonTg and 3xTg-AD mice aged 6, 12, 18 months, and in 18-month-old animals fed either a control diet or a high fat diet, no difference was observed in microvascular ACE2 levels. ACE2 levels in the mouse brain are not influenced by age or a diet that exacerbates AD-like neuropathology. Examples were taken from the same immunoblot experiment, and consecutive bands loaded in random order are shown. Actin and cyclophilin B are shown as loading controls. Data are represented as mean ± SEM. Statistical analysis: Kruskal-Wallis, ns, non-significant. *Abbreviations: Angiotensin-Converting Enzyme 2; Non-Tg,/NT, non-transgenic mice; 3x, 3xTg-AD mice; CD/C, control diet; CyB, cyclophilin B; HFD/H, high-fat diet; O.D., optical density*.

## Discussion

This present *postmortem* study investigated ACE2 concentrations in the brain of individuals with AD from two different cohorts. We assessed ACE2 protein levels in all subjects and mRNA expression in a subset. We observed a significant relationship between ACE2 levels, the neuropathological diagnosis of AD, and *antemortem* cognitive evaluation. Overall, our data indicate that (1) levels of TBS-soluble ACE2 in the parietal cortex were higher in persons with AD when compared to control subjects, accompanied by an elevation in ACE2 mRNA transcripts; (2) lower cognitive scores were associated with higher levels of ACE2 in TBS-soluble and cerebrovascular fractions; (3) an apparent transfer of ACE2 from membranes to soluble compartment was associated with pericyte loss and other markers of AD progression; (4) ACE2 levels remained unchanged in an animal model of AD-like neuropathology; (5) whereas ACE2 was concentrated in microvessels in the mouse brain, it was predominantly located in neurons in the human brain. Such a series of observations highlight that an AD diagnosis is associated with higher levels of specific forms of ACE2 in the brain, which might contribute to the higher risk of SARS-CoV-2 CNS infection in cognitively impaired individuals.

### Higher levels of soluble ACE2 are associated with AD and cognitive decline

The present observation of higher levels of soluble ACE2 in AD is in agreement with a previous report using a limited number of hippocampal samples of AD subjects (n = 13) compared to Controls (n = 5) [17]. Furthermore, preliminary human brain microarray data mentioned in a letter to the Editor also cite higher ACE2 expression levels in AD patient [47]. Although an association between SARS-CoV-2 infection and cognitive impairment has been previously evidenced at the population level [79] and hinted by genetic studies [25, 42], a significant correlation between ACE2 in the brain and cognitive scores has not been reported previously.

Several mechanisms could explain the higher levels of ACE2 in AD. Since old age increases the risk of infection with SARS-CoV-2 and developing cognitive decline and AD, we could have expected an association between cerebral ACE2 levels and advanced age. However, no correlation between age and ACE2 could be evidenced here in both human and mouse samples, suggesting that changes in TBS-soluble ACE2 are not directly related to age but rather to AD pathology and cognitive decline. This is consistent with health records data showing that dementia is associated with a higher risk for COVID-19, independently of age [79]. Indeed, a recent network analysis suggest that AD and COVID-19 share defects in neuroinflammation and microvascular injury pathways [90]. Second, the increase in ACE2 could be a consequence of the AD neurodegenerative process and/or of potential compensatory mechanisms in response to AD neuropathology, which could include an increase in gene transcription, which is consistent with the higher mRNA expression measured in AD samples. However, data gathered in 3xTg-AD mice do not fully support that hypothesis. Indeed, the accumulation of human Aβ peptides and hyperphosphorylated tau in this mouse model did not lead to changes in murine ACE2 protein levels, suggesting that an ACE2 increase is not a consequence of classical AD neuropathology. Nevertheless, it should be reminded that such a mouse model displays an amount of Aβ and tau 1 to 3 logs lower than what is typically found in an AD brain. Moreover, ACE2 pathways may simply be regulated differently in the mouse, as suggested by the different localization of the protein in the murine brain.

ACE2 is part of the renin angiotensin system (RAS), which regulates the vascular system. An increase of cerebral ACE2 may impact the brain RAS, thereby affecting blood flow, arterial pressure, neuroinflammation and, consequently, brain function. Such a dysregulation of the RAS- equilibrium in the brain could contribute to the aetiology of several neurodegenerative diseases, including AD [2, 80, 81]. For example, in a cohort study including community-dwelling older adults with mild to moderate AD, the use of ACE inhibitors (ACEi) was associated with a slower cognitive decline, independent from their antihypertensive effects [69]. ACEi and angiotensin II receptor blockers (ARBs) are also under investigation to improve cognitive impairment associated with AD [24, 27, 37, 65]. Moreover, twice higher soluble ACE2 has been reported in the cerebrospinal fluid of hypertensive patients [83]. In our study, we did not detect any association with the use of drugs acting on ACE, such as ARBS or ACEi, and brain levels of ACE2, but the study was not designed for that purpose. However, it is important to note that the levels of ACE2 detected by immunoblotting may not directly inform on ACE2 activity. Indeed, a *postmortem* assessment of ACE2 enzymatic activity with a fluorogenic assay instead showed a reduction in AD [43]. Studies in animals indicate that pharmacological activation of ACE2 rather reduces hippocampal soluble Aβ and reverses cognitive impairment in the Tg2576 model of Aβ neuropathology [22].

Another peculiar observation is the difference between ACE2 found in soluble fractions containing cytoplasmic/extracellular proteins *versus* ACE2 retrieved in detergent-soluble fractions containing membrane-bound proteins. Overall, ACE2 in TBS-soluble fractions was higher in subjects with AD, while no such trend was observed with ACE2 located in membranes. Moreover, the correlation between ACE2 and AD-relevant markers, most notably the pericyte markers PDGFRβ and others like ANPEP, differed significantly between the two fractions. The strong inverse association with TDP-43 pathology was also limited to detergent-soluble ACE2. Previous studies did not distinguish TBS-soluble *versus* detergent-soluble ACE2 [17, 47]. Although ACE2 is generally considered a membrane protein, its actual attachment to the cytoplasm membrane is relatively weak. For example, the ACE2 ectodomain can be cleaved by ADAM17 or TMPRSS2 and released in the cytoplasm [35, 91]. Recent studies report that a decrease in active membrane-bound ACE2 due to ADAM17 and TMPRSS2 overactivation could be deleterious for SARS-CoV-2-infected patients [35, 60, 82]. However, we did not observe differences in mRNA and protein levels of TMPRSS2. Vascular ACE2 was also specifically measured in this study. Despite associations with cognitive scores and PDGFRβ levels, no significant difference was detected between groups, possibly due to the interindividual variability induced by the separation process. An intriguing possibility explaining the higher content in ACE2 specific in the TBS fraction, as detected with an antibody targeting the N-terminal extracellular domain, could be an enhanced release of ACE2 from the membrane to the cytosol or the extracellular parenchyma in AD, also termed ACE2 shedding [35, 78]. Such a detachment of ACE2 from cell membranes may be a pathological phenomenon associated with AD, warranting further study.

The present work also unveils additional information on the cellular localization of ACE2 in the human brain. While we observed an enrichment of ACE2 in mouse microvessels, such was not the case in human samples. ACE2 was detected in human samples using Western blot and RT-qPCR, but a clear immunofluorescence signal was more challenging to achieve. Unlike in the mouse, where the enrichment in microvessels was evident, the detection of ACE2 in human brain capillaries became apparent only after microvascular fractionation. However, ACE2 was clearly present in neurons in human brain sections, corroborating Western blot results. Nonetheless, it should be noted that brains from mice were harvested quickly after transcardial perfusion. On the other hand, human brain tissue underwent *premortem* and *postmortem* events, which may have affected ACE2 distribution and detection. In sum, the present data obtained using several different antibodies indicate that the cerebral distribution of ACE2 is less strictly vascular, more neuronal in humans compared to the mouse. At the very least, work related to human ACE2 but performed in mouse models should be interpreted with caution regarding their possible application to the brain RAS, AD and other neuropathologies, as well as central SARS-CoV-2 infection in humans.

## Conclusions

In summary, the present data show an accumulation of the soluble form of ACE2 associated with cognitive decline in individuals with a neuropathological diagnosis of AD. ACE2 levels were not influenced by age or biological sex. We also observed a strong association between soluble ACE2 levels and AD neuropathology, as well as pericyte loss. The search for molecular cues that cause a rise of TBS-soluble ACE2 and regulate the brain RAS in AD subjects may ultimately lead to the discovery of new therapeutics to prevent cognitive decline and AD.

## List of Abbreviations

ACEi: Angiotension I Converting Enzyme Inhibitors
ACE2: Angiotensin I Converting Enzyme 2
AD: Alzheimer’s disease
ARBs: Angiotensin II receptors blockers
BA: Brodmann area
BBB: Blood brain barrier
CD: control diet
CNS: Central Nervous System
COVID-19: Coronavirus disease 2019
DAPI: 4′,6-diamidino-2-phenylindole
EHR: electronic health records
FFPE: formalin-fixed, paraffin-embedded
GWAS: genome wide associations study
HRP: horseradish peroxidase
HFD: High fat diet
MCI: Mild-cognitive impairment
NIA-AA: National Institute of Aging – Alzheimer’s Association
NCI: No cognitive impairment
O.D.: Optical density
RBD: Receptor Binding Domain
NHS: Normal Horse Serum
PBS: phosphate-buffered saline
pCAA: parenchymal cerebral angiopathy amyloid
PDGFRβ: Platelet-derived growth factor receptor β
ROS: Religious Order Study
SARS-CoV-2: Severe Acute Respiratory Syndrome CoronaVirus 2
SDS-PAGE: sodium dodecyl sulphate-polyacrylamide gel electrophoresis
TBS: Tris-Buffered Saline
TMPRSS2: Transmembrane protease serine 2.

## Declarations

### Ethics approval and consent to participate

All procedures performed with volunteers included in this study were in accordance with the ethical standards of the institutional ethics committees and with the 1964 Helsinki Declaration. Written informed consent was obtained from all individual participants included in this study. All procedures relating to mouse care and experimental treatments were approved by the Laval University animal research committee (CPAUL) in accordance with the standards of the Canadian Council on Animal Care.

### Consent for publication

Not applicable.

### Availability of data and material

The datasets analysed during the current study available from the corresponding author on reasonable request. Data from the ROS can be requested at https://www.radc.rush.edu.

### Authors’ contributions

LR, VE, SH, and FC designed the study. LR, ML, VE, AL, PB, and CT performed experiments. DAB provided ROS samples. LR and FC analyzed data. LR, VE and FC wrote the first drafts of the paper.

## Acknowledgements

We thank Dr Pierre Leclerc from CRCHU de Québec – Université Laval for providing human FFPE testis samples. The authors are indebted to the nuns, priests and brothers from the Catholic clergy participating in the Religious Orders Study.

## Supplementary material

Supplementary material is available online.

---

**Figure S1:**
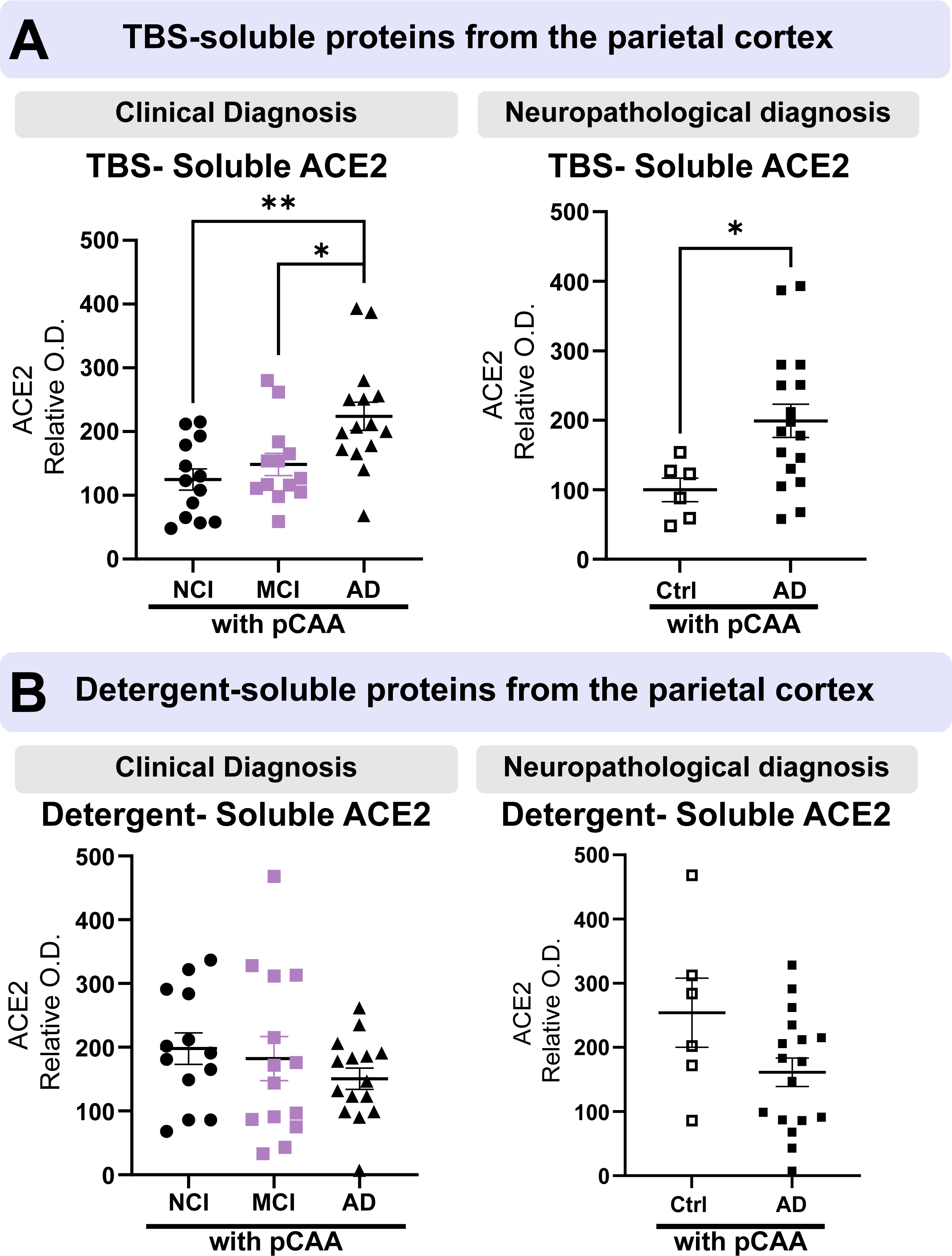
AD subjects with parenchymal CAA have higher soluble ACE2. Individuals were grouped based on their ABC neuropathological diagnosis or clinical diagnosis and subdivided based on the presence of parenchymal CAA (pCAA). *(A)* Levels of soluble ACE2 were higher in subjects with an AD neuropathological and clinical diagnosis with the presence of pCAA. *(B)* Levels of detergent-soluble ACE2 did not change depending on the presence or absence of pCAA. All samples, loaded in a random order, were run on the same immunoblot experiment for quantification. Examples were taken from the same immunoblot experiment, and consecutive bands loaded in random order are shown. Data are represented as a scatterplot. Horizontal lines indicate mean ± SEM. Statistical analysis: two groups: unpaired t-test *p <0.05 or three groups: Tukey’s multiple comparisons test *p<0.05, **p<0.01. *Abbreviations: A/AD, Alzheimer’s disease;; C, control; Clinical Dx, clinical diagnosis; M/MCI, mild cognitive impairment; N/NCI, healthy controls with no cognitive impairment; O.D., Optical density; pCAA, parenchymal cerebral amyloid angiopathy*.

**Figure S2:**
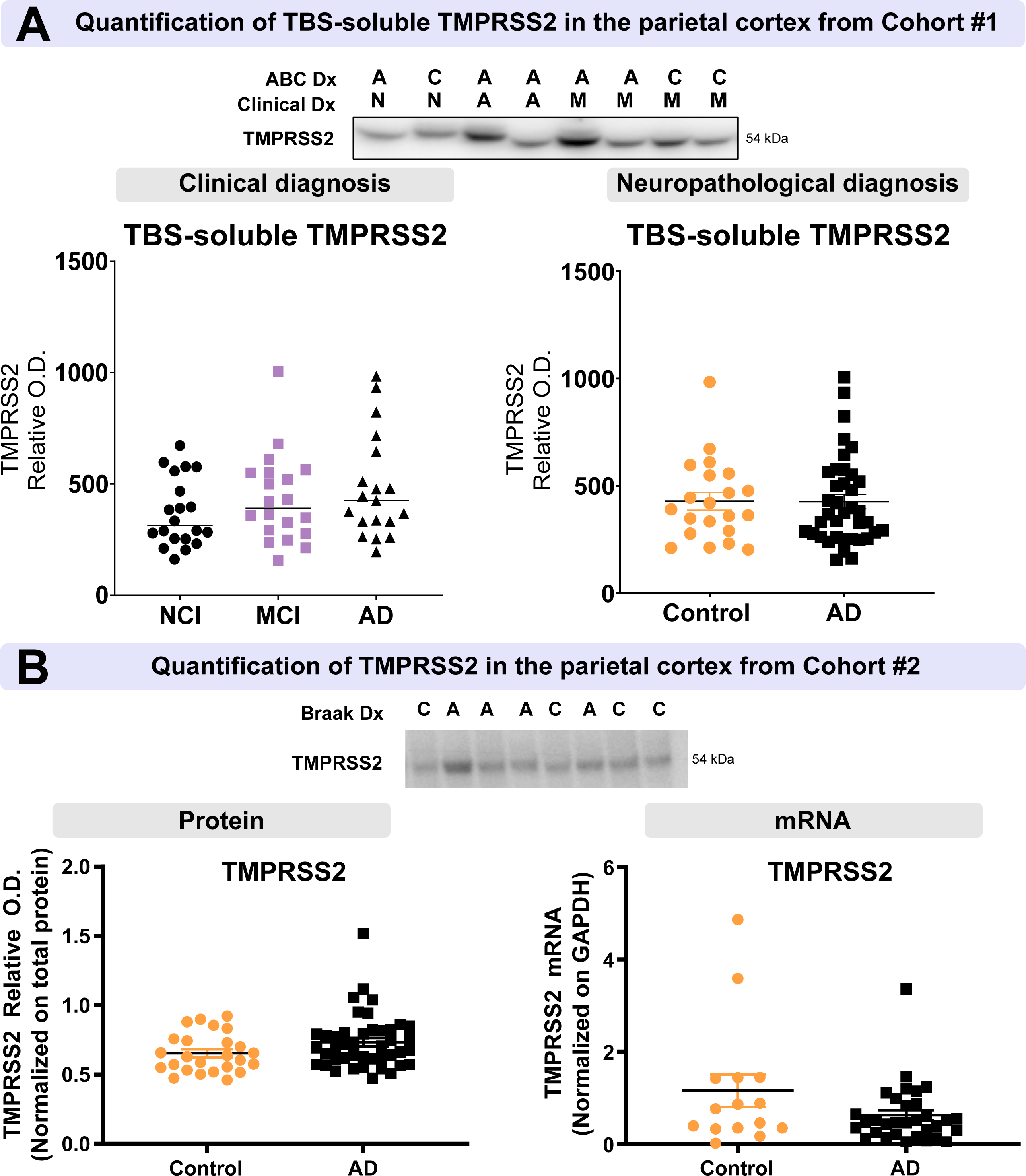
Levels of TMPRSS2 protein, which is employed by SARS-CoV-2 for Spike protein priming, are unchanged in AD individuals. *(A)* Levels of the TMPRSS2 protein were measured in Cohort #1: no difference was identified according to clinical or neuropathological (ABC) diagnosis. *(B)* In Cohort#2, TMRPSS protein and mRNA quantification did not reveal difference between control and AD. All samples, loaded in a random order, were run on the same immunoblot experiment for quantification. Examples were taken from the same immunoblot experiment, and consecutive bands loaded in random order are shown. Data are represented as a scatterplot. Horizontal lines indicate mean ± SEM. Statistical analysis: Ordinary on-way ANOVA or Mann-Whitney test, non-significant. *Abbreviations: A/AD, Alzheimer’s disease; ABC Dx, ABC neuropathological diagnosis; Braak Dx, Braak staging diagnosis; C, control; Clinical Dx, clinical diagnosis; M/MCI, mild cognitive impairment; N/NCI, healthy controls with no cognitive impairment; ROD, relative optical density; TMRPSS2, Transmembrane protease serine 2*.

**Figure S3:**
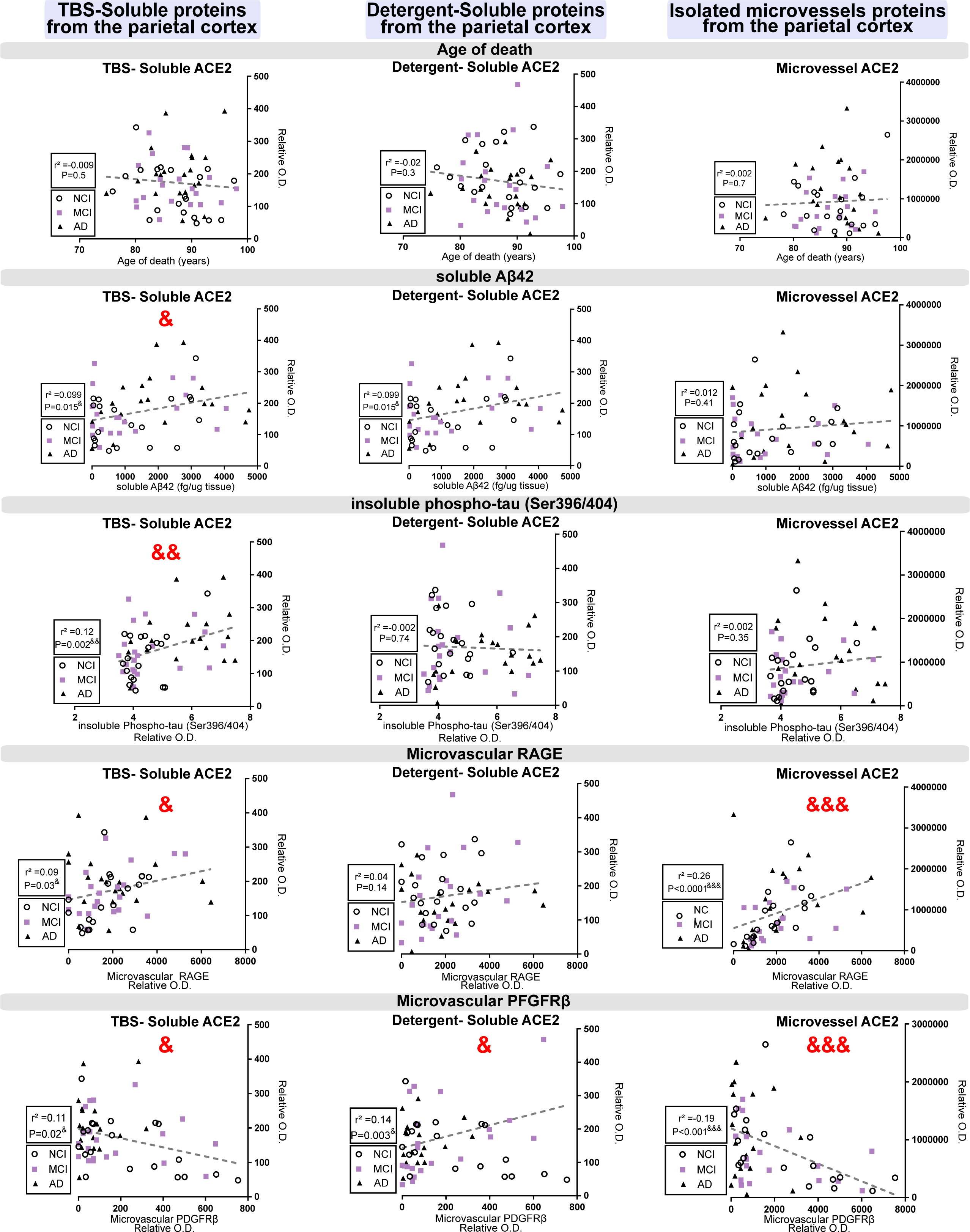
Examples of correlation plots between ACE2 in the three protein fractions and age or AD-related proteins are shown. No significant correlation was established between ACE2 in all fractions tested and the age of death. TBS-soluble ACE2 was positively correlated with soluble Aß42 peptides and phospho-tau AD2. Both TBS-soluble and microvessel ACE2 were positively associated with microvascular RAGE but negatively with microvascular PDGFRß. Significant correlations are shown with the sign & in red. All proteins presented in these correlations were determined by Western-blot analysis except Aß peptides, which were determined using ELISA. Coefficient of determination &p < 0.05, &&p < 0.01, &&&p < 0.001. *Abbreviations: ACE2, Angiotensin-Converting Enzyme 2; AD, Alzheimer’s disease; MCI, mild cognitive impairment; NCI, healthy controls with no cognitive impairment; PDGFRß, Platelet Derived Growth Factor Receptor Beta; RAGE, Receptor for Advanced Glycation Endproducts; Relative OD, relative optical density*.

**Figure S4:**
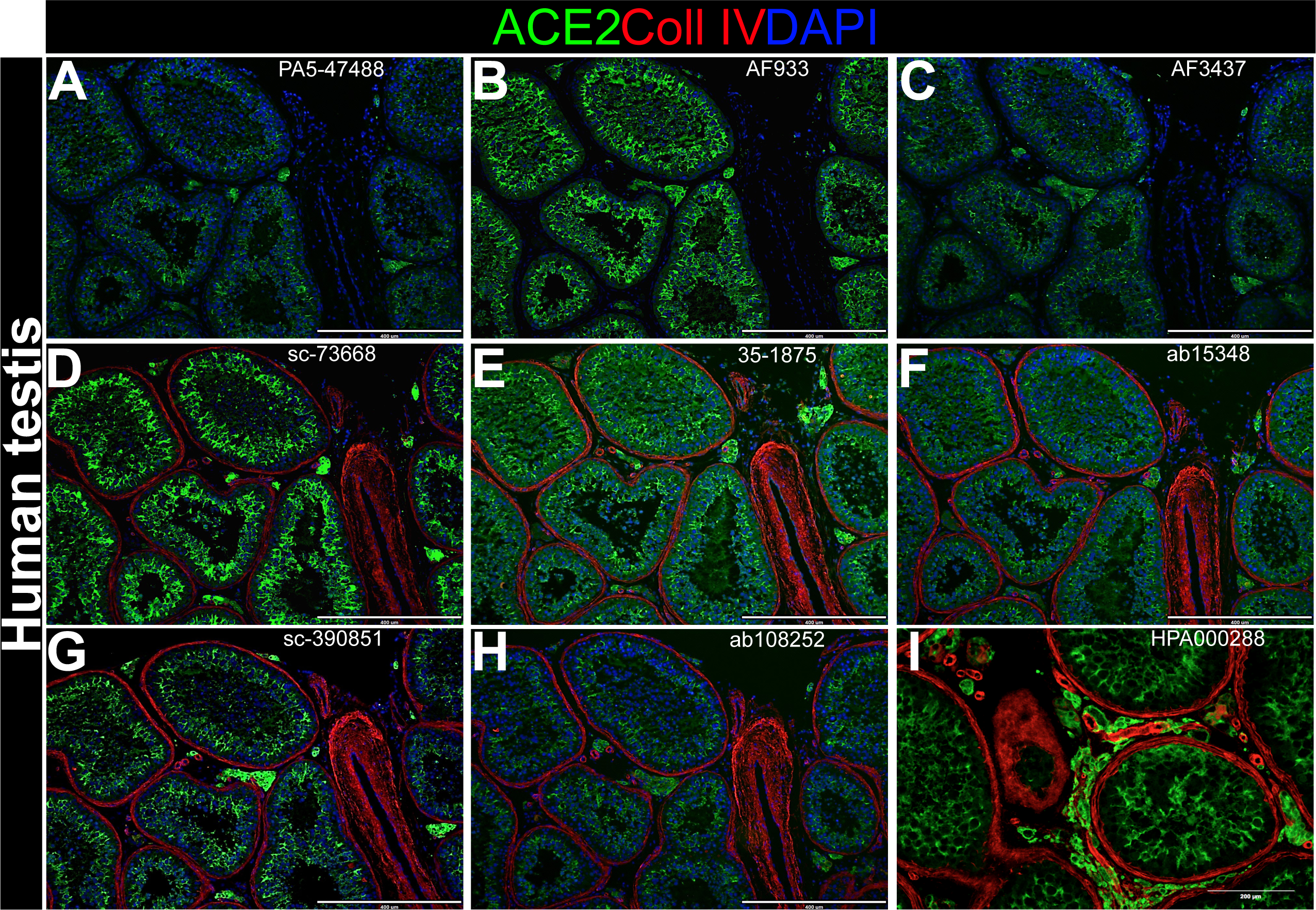
Human formalin-fixed paraffin-embedded testis sections were used as positive controls for ACE2 immunostaining, which is strong in this tissue. Antibodies used: (A) goat pAb #PA5-4788 from ThermoFisher, (B) goat pAb #AF933 from BioTechne, (C) goat pAb #AF3437 from BioTechne, (D) mouse mAb #sc-73668 from Santa Cruz, (E) rabbit pAb #35-1875 from Abeomics, (F) rb pAB #ab15348 from Abcam, (G) mouse mAb #sc-390851 from Santa Cruz, (H) rabbit mAb #ab108252 from Abcam and (I) rabbit pAb #HPA000288 from Atlas. Scale bars: 400 µm (A-H) and 200 µm (I). *Abbreviations: ACE2, Angiotensin-Converting Enzyme 2; Coll IV, Collagen IV; mAb, monoclonal antibody; pAb, polyclonal antibody*.

**Figure S5:**
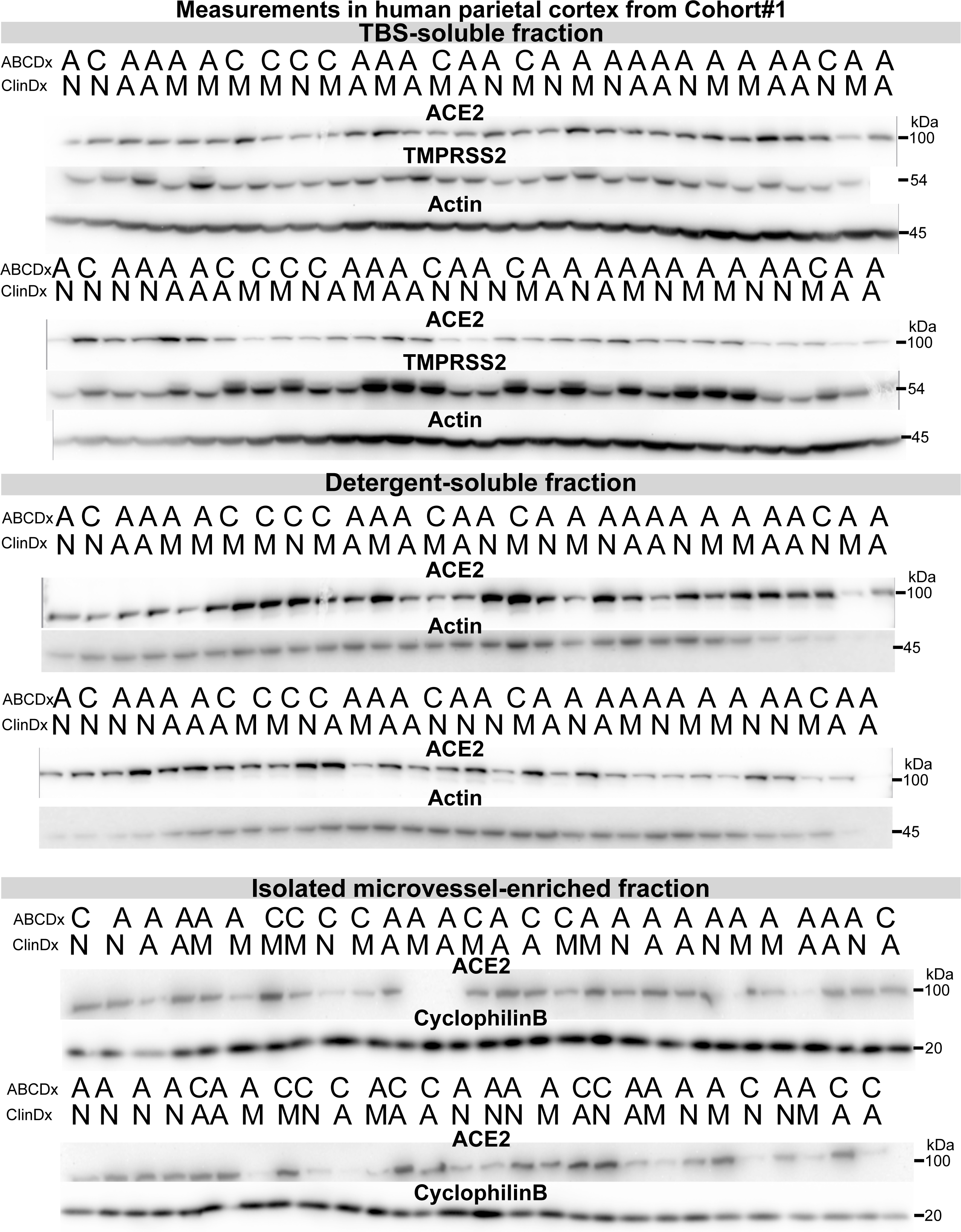
Western blots in human TBS-soluble, detergent-soluble and microvessel-enriched extracts from Cohort#1. The clinical and neuropathological diagnoses are given above each sample. *Abbreviations: A, Alzheimer’s Disease; ABC Dx, ABC neuropathological diagnosis; ACE2, Angiotensin-Converting Enzyme 2; C, Control; ClinDx, Clinical diagnosis; M, mild cognitive impairment; N, healthy controls with no cognitive impairment; TMRPSS2, Transmembrane protease serine 2*.

**Figure S6:**
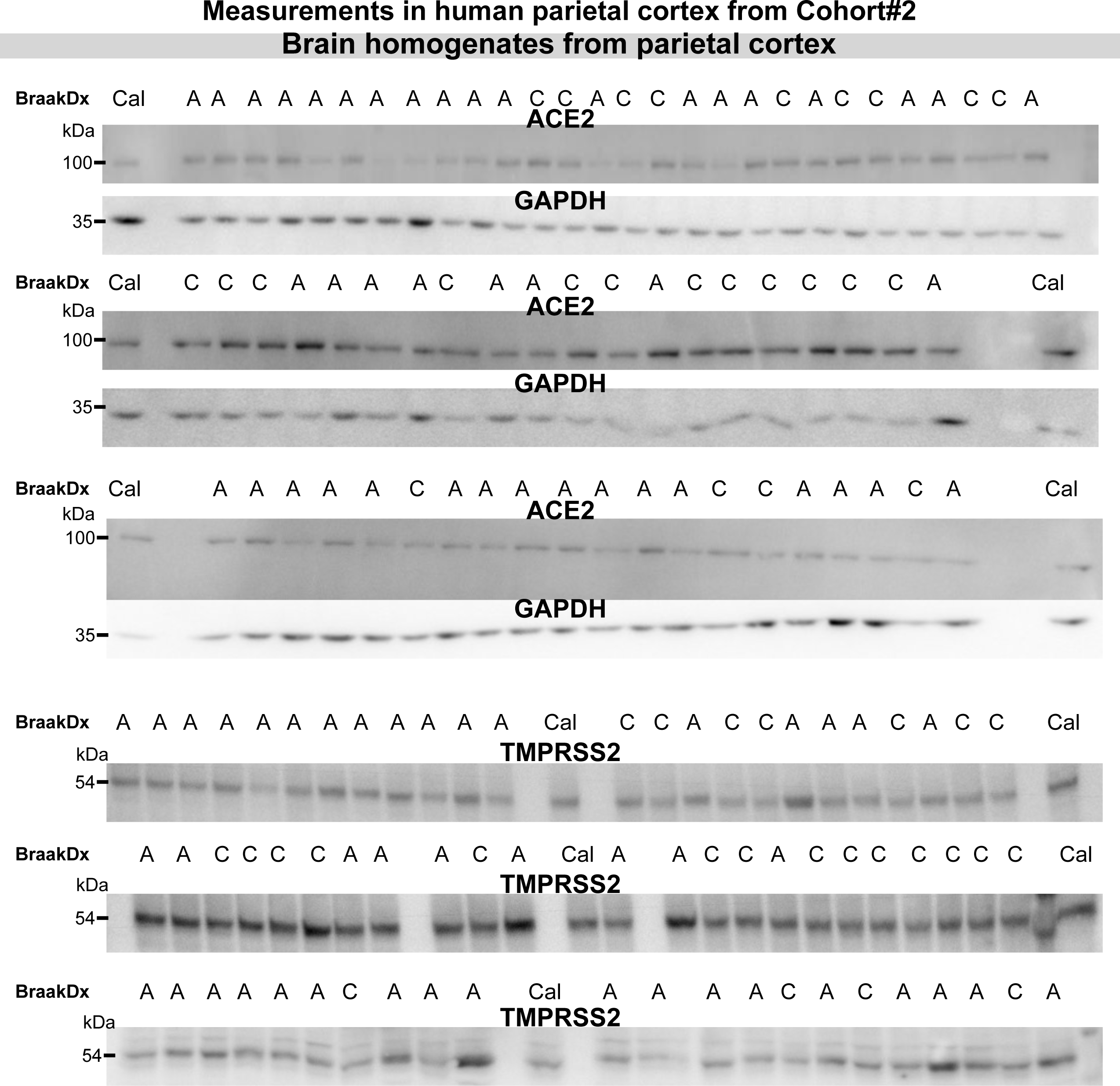
Western blots in brain homogenates from Cohort#2. Diagnosis was determined using Braak staging. Abbreviations: A, Alzheimer’s Disease; ACE2, Angiotensin-Converting Enzyme 2; Braak Dx, Braak staging diagnosis; Cal, Calibrator; C, Control; GAPDH, Glyceraldehyde 3-phosphate dehydrogenase; TMRPSS2, Transmembrane protease serine 2.

## Supplementary method

### Isolation of murine brain microvessels

The procedure used for isolation of murine brain microvessels has been reported in our previous work (Bourassa *et al*., 2019a). Nontransgenic and 3xTg-AD mice aged 6, 12 and 18 months were sacrificed with an intracardiac perfusion of ice-cold PBS containing 0.32 M sucrose and protease (SIGMA*FAST* Protease Inhibitor tablets, Sigma-Aldrich) and phosphatase (1 mM sodium pyrophosphate and 50 mM sodium fluoride) inhibitors, under deep anesthesia with ketamine/xylazine. The brains were immediately collected, and brainstem, cerebellum and meninges were removed. Murine brain samples were then chopped and frozen in 0.5 mL of MIB containing 0.32 M sucrose and protease and phosphatase inhibitors (Bimake). For a milder freezing we used Mr. Frosty™ Freezing Container (Thermo Scientific). The microvessel enrichment procedure was then conducted as described for human samples. To validate the enrichment of mural cell markers, the microvessel-enriched and the microvessel-depleted fractions were compared to a total brain homogenate obtained from the homogenization of a whole hemisphere of a control mouse in the lysis buffer. Protein concentrations in all fractions were determined using the bicinchoninic acid assay (Thermo Fisher Scientific)

